# Distinct transcriptomic and epigenomic responses of mature oligodendrocytes during disease progression in a mouse model of multiple sclerosis

**DOI:** 10.1101/2023.12.18.572120

**Authors:** Chao Zheng, Bastien Hervé, Mandy Meijer, Leslie Ann Rubio Rodríguez-Kirby, André Ortlieb Guerreiro Cacais, Petra Kukanja, Mukund Kabbe, Tomas Olsson, Eneritz Agirre, Gonçalo Castelo-Branco

## Abstract

Multiple sclerosis (**MS**) is a chronic demyelinating autoimmune disease that targets mature oligodendrocytes **(MOLs**) and their myelin. MOLs are transcriptionally heterogeneous and can transition to immune-like states in the context of MS. However, the intricacies of their dynamics throughout disease progression remain poorly understood. Here, we employed simultaneous single-cell multiome ATAC and RNA sequencing targeting oligodendroglia (**OLGs**) from the experimental autoimmune encephalomyelitis (**EAE**) MS mouse model at different stages of the disease course. We found that the transition to immune OLG states appear already at the early stages of EAE and persist to the late stages of the disease. Interestingly, transcription factor activity suggested immunosuppression in MOLs at early stages of EAE and we also observed a transitory activation of a regenerative program in MOLs at this stage. Importantly, different MOLs exhibit a differential responsiveness to EAE, with MOL2 exhibiting a stronger transcriptional immune response than MOL5/6. Moreover, we observed divergent responses at the epigenetic level of MOL2 and MOL5/6 during disease evolution. Thus, our single-cell multiomic resource highlights dynamic and distinct responses of OLG subpopulations to the evolving environment in EAE, which might modulate their response to regenerative therapeutic interventions in MS.

## Introduction

Multiple sclerosis (**MS**) is an inflammatory autoimmune disease of the central nervous system (**CNS**) which typically develops at a young age, in the 20s-30s. There are currently four accepted disease courses/stages of MS: clinically isolated syndrome (**CIS**), relapsing-remitting MS (**RRMS**), primary progressive MS (**PPMS**), and secondary progressive MS (**SPMS**)^1^. Some of the pathological characteristics of MS are the infiltration of immune cells and mature oligodendrocytes (**MOLs**) impairment, which result in demyelinated lesions in the CNS and ultimately lead to physical disability, cognitive impairment, and a decrease in life quality. The symptoms and annualized relapse rate of MS patients can be reduced with the application of disease-modifying therapies. Nevertheless, the therapeutic effect varies among individuals and these treatments are not particularly effective for PPMS and SPMS^2^.

Oligodendrocytes (**OLs**) are the myelinating cells of the CNS, which provide support to the axons of neurons and ensure fast saltatory conduction of action potentials. Oligodendrocyte precursor cells (**OPCs**), which originate in the ventricular zones of the embryonic neural tube, are present throughout the developing and adult CNS, and are capable of differentiating into myelinating MOLs^3^. In the pathogenesis of MS, lymphocytes-mediated autoimmune responses are activated and specifically affect myelin sheaths and MOLs. MS has been generally viewed as primarily driven by T cells and B cells^4^. However, recent studies have revealed the expression of immunomodulatory molecules in oligodendroglia (**OLG**) not only in MS, but also in AD and aging^5–9^. OPCs expressing MHC-I and -II also have been shown to process and present antigens, resulting in the activation of CD8+ cytototix T cells (CTLs^)6^ and CD4+^5^. These findings indicate the potential role of oligodendroglia in the modulation of immune responses within the CNS.

Single-cell/nucleus RNA-sequencing (**scRNA-seq**) is a powerful technique for analyzing transitions between cellular states in homeostasis and disease. It has been applied to OLGs from human and mouse samples, and it has revealed specific MOLs/OPC sub-populations in MS and EAE^5, 10^. However, these studies were conducted with samples from single time points, at the peak of the disease in mouse EAE, and from the late stage of the disease (in human post-mortem MS tissue material). Thus, the dynamics of OLGs throughout the disease process have not been explored. In particular, it is unclear when the transition of OLGs to immune-like states occurs. Given that MS is a disease with profound heterogeneity in the immunopathological patterns between different disease stages, it is highly pertinent to determine changes in the cellular states of OLGs during the time course of the disease.

Reshaping of transcriptional and epigenetic landscapes underlie transitions between distinct cellular states. Assay for Transposase-Accessible Chromatin using sequencing (**ATAC-seq**) is a high throughput sequencing technique for assessing genome-wide chromatin accessibility by using hyperactive Tn5 transposase to insert sequencing adapters into accessible chromatin regions^11^. ATAC-seq provides information on regions with open chromatin, which is required for gene expression by allowing the binding of the RNA polymerase II machinery, transcription factors and other regulatory proteins. Enhancer and promoter regions efficiently regulate gene expression by binding to transcription factors. ATAC-seq can also be used in combination with other genomic techniques, such as RNA-seq, to study the dynamics of gene expression at the transcriptome level and its correlation with chromatin accessibility^11^. By applying scATAC-seq alone and in combination with scRNA-seq, we previously found that a cohort of immune genes exhibit open chromatin in both control and EAE at peak disease, suggesting a primed chromatin state, while their expression only increases in the context of EAE^12^.

Here, we investigate the epigenomic and transcriptional dynamics of OLG in the course of EAE disease evolution. We applied simultaneous single-cell multiome ATAC and RNA sequencing to OLGs FACS-sorted from male and female EAE at three distinct time points:1) early stage, when neurological symptoms begin to arise; 2) peak, after symptoms rapidly increased; 3) late stages, when the disease is chronic and progression has stabilized. Strikingly, at the early stage of EAE a subset of genes involved in antigen presentation showed increased expression and chromatin accessibility in MOLs, indicating that the induction of an immune-like state in OLG occurs before the formation of fully developed lesions. Moreover, chromatin accessibility at these genes remained highly open at the late stage of EAE, indicating either a partial maintenance or an epigenetic memory of this immune-like state. We also found that cholesterol biosynthesis processing genes showed a transitory increase in MOLs at the early stage, indicating a direct regenerative response to the first signs of inflammation in the CNS. Furthermore, specific MOL subtypes acquire different patterns of change in genes related to immune response and OLG differentiation at both expression and chromatin accessibility levels during the course of disease. We observed that the white matter-enriched^13^ MOL2 showed higher immune signatures than MOL5/6, which in turn exhibited induction of chromatin accessibility in genes with regenerative pathways, particularly in the late stage. Thus, different MOLs populations might react and contribute to disease evolution in distinct manners. Our study provides a unique resource (available for browsing at the UCSC Cell Browser and Genome Browser (https://olg-dyn-eae-multiome.cells.ucsc.edu) and a deeper understanding of the OLG epigenomic and transcriptomic dynamics in the inflammatory demyelination mouse model of MS, possibly allowing better therapeutic targets for achieving immune modulation and myelin regeneration in MS.

## Results

### Simultaneous chromatin accessibility and transcriptomic single-cell analysis of OLG at different stages of EAE

We induced EAE in *Sox10:Cre-RCE:LoxP* (EGFP) transgenic mice^13^ lineage-trace OLG, with injection of emulsion containing the MOG 35-55 immunogenic peptide in complete Freund’s adjuvant (CFA), followed by intraperitoneal injection of pertussis toxin. Spinal cord tissues from male and female mice induced with EAE were collected at three different stages: 1) early stage (day 8-9 post-injection, score 0-0.5, 8 mice in 4 multiome experiments); 2) peak stage (day 14-15, score 3, 8 mice in 4 multiome experiments); 3) late/chronic stage (day 37-40, score 2-2.5, 10 mice in 5 multiome experiments) (**Fig. 1a,b**). Spinal cord samples from CFA-Ctrl were also collected from the same stages (early stage, peak stage, late stage) with a score of 0 (4 mice in 2 multiome experiments for each stage), alongside spinal cords tissues from non-induced Naïve-Ctrls (3-month-old mice from the same strain, 4 mice in 2 multiome experiments). OLG were enriched based on EGFP by FACS sorting (**Extended Data Fig. 1a**), after which we performed single cell multiome RNA-seq and ATAC-seq (**Fig. 1a**).

**Fig. 1.**
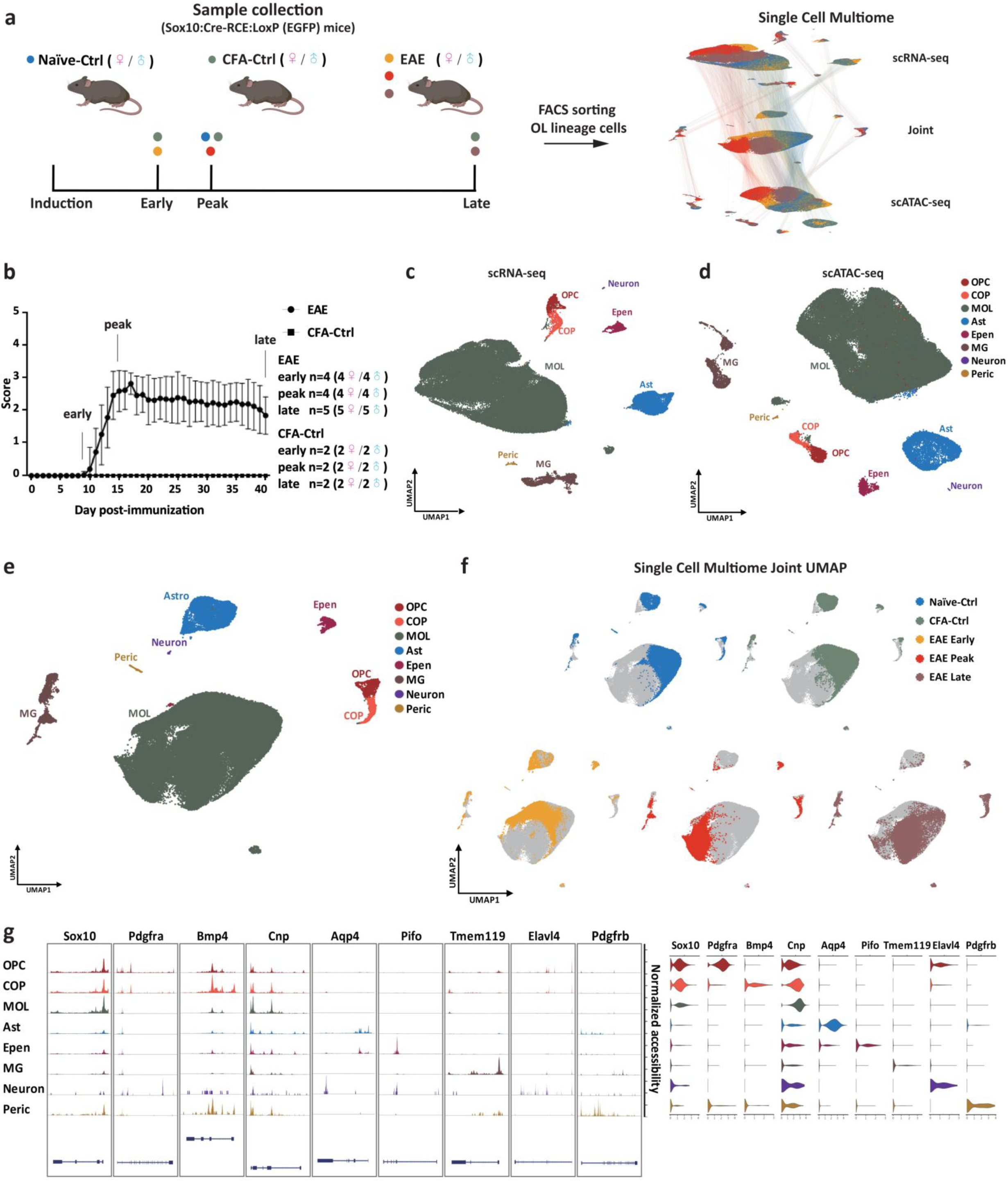
Single cell multiome in oligodendroglia (OLGs) from the EAE mouse model of MS showing transition between transcriptional/epigenomic states among different time points. **a**, Schematic of the methodology used in animal model establishment and multiome sequencing (created with BioRender.com). Dots in blue: Naïve-Ctrls, green: CFA-Ctrl, yellow: EAE early stage, red: EAE peak stage, brown: EAE late stage. **b**, Clinical score of the mice used in the study (EAE: 26 mice in 13 multiome experiments, CFA-ctrl n = 6; data represented as mean ± SD). For EAE from peak and late stages, only mice that had reached score 3 were used in this study. **c, d**, Uniform manifold approximation and projection (UMAP) of cells profiled with simultaneous scRNA-seq (c) and scATAC-seq (d). Cell types are identified according to marker genes. Ast, astrocytes; EP, ependymal cells; MG, microglia; MOL, mature oligodendrocytes, OPC, oligodendrocyte progenitor cells. **e**, Joint UMAP from the weighted nearest neighbors graph of scRNA-seq and scATAC-seq modalities. **f**, Joint UMAP with cells separated by conditions, on top of UMAP with all cells (color in gray). **g**, Normalized chromatin accessibility (left) and Iog2 expression (right) of representative marker genes in each cell type.

After sample-specific quality-control filtering (**Extended Data Fig. 1b, see methods**), we obtained 91,757 cells in total (Naïve-Ctrl 12,721 cells, CFA-Ctrl 25,932 cells, EAE early stage 12,808 cells, EAE peak stage 14,255 cells, EAE late stage 26,041 cells). Clustering with the Louvain algorithm identified 35 clusters based on the scRNA-seq dataset (**Extended Data Fig. 1c,e**), and 33 clusters based on the scATAC-seq dataset (**Extended Data Fig. 1d,f**). Since the scRNA-seq and scATAC-seq data were derived from the same cell, we used the reduced dimensions from both modalities to generate a new joined projection (**Fig. 1e,f and Extended Data Fig. 1g,h**), resulting in 31 clusters, annotated by cell type with marker gene expression and chromatin accessibility (**Fig. 1g**, see Methods). As expected from the lineage tracing strategy, the majority of the cells were MOLs, OPCs, and committed oligodendrocyte precursors (**COP**). Nevertheless, other populations, in particular astrocytes (**Ast**) and microglia (**MG**), were also captured (**Fig. 1c-e**).

### Transition to immune OLG states occurs in early stages of EAE and persists to late stages

OPCs and MOLs have been previously shown to acquire disease-associated states at the peak of EAE^12^. Multiome analysis of MOLs at the peak stage indeed showed that they transitioned to transcriptional/epigenomic states distinct from CFA-Ctrl and any other time point (**Fig. 1f and Extended Data Fig. 1e,f**). Importantly, we could observe the onset of this transition for a subset of MOLs in the early stages of EAE, when lesions are just starting to develop (**Fig. 1f and Extended Data Fig. 1e,f**). Moreover, a minority of MOLs retained peak-stage-like profiles at the late stages, while most MOLs transitioned to CFA-Ctrl-like states, suggesting a return to homeostasis or transition to a new homeostatic state (**Fig. 1f and Extended Data Fig. 1e,f**).

Disease-associated OPC and MOL states are characterized by chromatin accessibility and expression of immune genes^5, 12^. Thus, we investigated whether the observed transitions were driven by the immune status of MOLs. For this purpose, we subsetted and reclustered OPCs and MOLs from CFA-Ctrl and the three EAE time points, to identify those expressing immune-related genes^12^(see Methods) (**Fig. 2a**). Immune oligodendroglia (**imOLGs**) were hardly observed in CFA-Ctrl spinal cords, but were found at early, peak and late stages, with a higher percentage present at the peak stage compared to the early and late stages (**Fig. 2c**, early stage 28.56%, peak stage 72.43%, and late stage 37.38%). Chromatin accessibility at the promoter/gene body of the same immune genes identified a lower number of imOLGs than by immune gene expression (early 13.29%, peak stage 31.49%, and late stage 15.92%) (**Fig. 2b,d**). This is most likely due to some OLGs from CFA and Naïve Ctrls exhibiting already primed chromatin accessibility in immune gene loci but with low or no expression^8,12^, which made the threshold of immune gene chromatin accessibility higher than the expression. Thus, our data indicates that the transition of OLGs to immune-like states at epigenomic and transcriptional levels occurs at early stages of EAE, when lesions are just starting to develop, and persists at late stages, despite resolving inflammation.

**Fig. 2.**
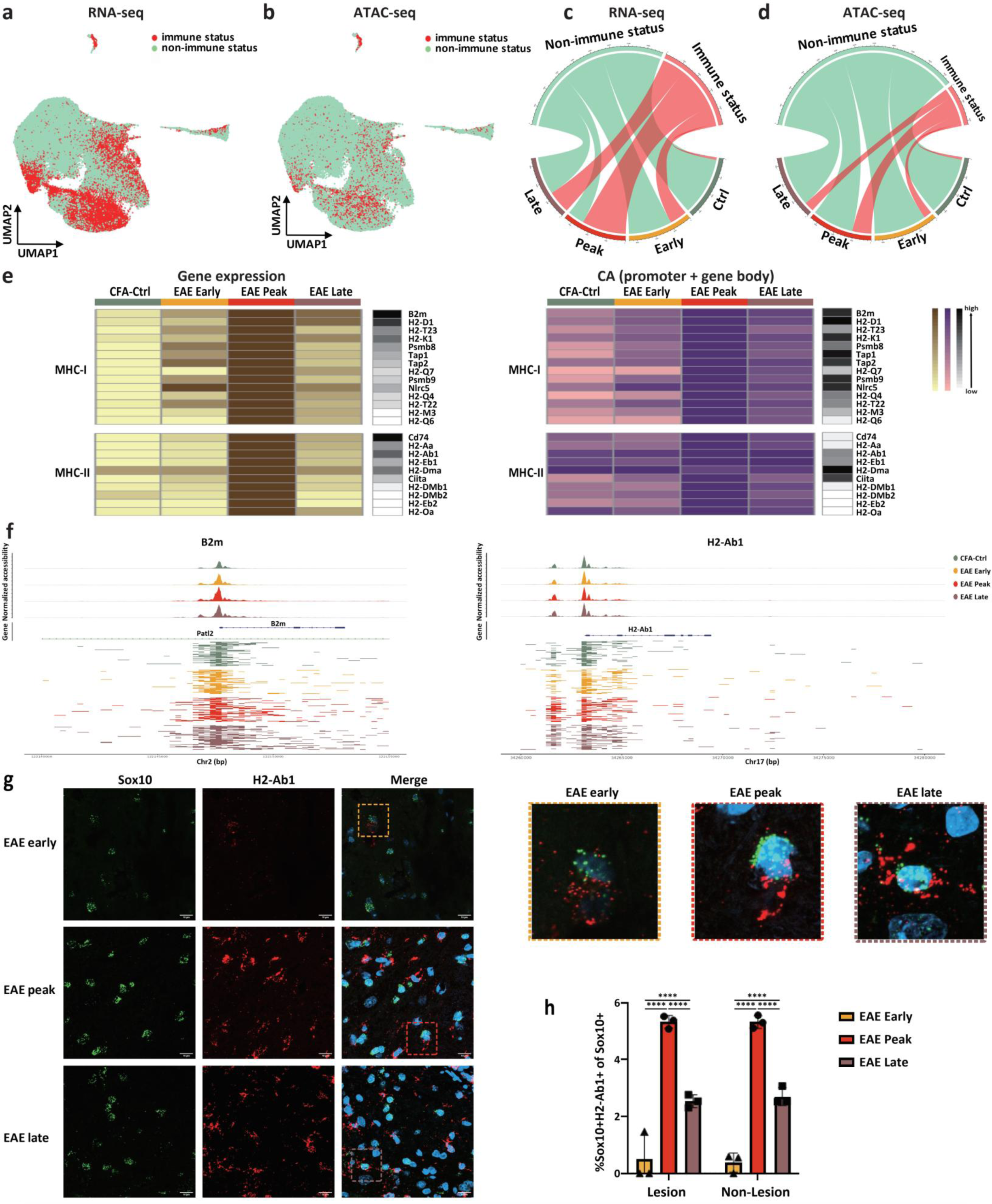
Transition to immune oligodendroglia states occurs already at early stages of EAE and persists at late stages. **a, b**, Joint UMAP with oligodendroglia from CFA-Ctrl and EAE. Projection of cells with immune status (red) and non-immune status (green) identified with gene expression (a) and chromatin accessibility (b). **c**,d, Number of cells with or without immune status for CFA-Ctrl and EAE identified with gene expression (c) and chromatin accessibility (d) at each time point (downsampled by time point). **e**, Heatmaps of the expression (left) and chromatin accessibility (CA, right) of MHC-I and -II genes at different stages. The black column on the side represents the gene raw counts. **f**, Normalized chromatin accessibility of representative MHC-I and MHC-II genes at different time points (top) and ATAC-seq reads alignment of 50 individual random cells per condition for the same genes (bottom). **g**, RNAscope ISH representing spinal cord sections from early, peak, and late stage EAE mice marked with probes for SoxlO (OLGs) and H2-Abl (MHC-II). **h**, Quantification of the percentages of Soxl0+H2-Abl+ cells out of Soxl0+ cells in lesion and non-lesion areas. **** P<O.OOOl; n = 3 independent experiments per condition; data represented as mean ± SD.

### Persistence of chromatin accessibility at MHC-I and -II genes in OLG at late stages of EAE

The major histocompatibility complex (**MHC**) gene clusters encode proteins involved the activation of T cells. The expression and chromatin accessibility of some MHC-I and -II related genes, such as *Psmb9*, *Tap1*, and *H2-Ab1*, were previously shown to be increased in EAE-specific OLGs at peak stages of EAE^5, 12^. We thus examined the expression and chromatin accessibility at the promoter/gene body of MHC-I and -II genes in OLGs across different EAE stages. We found that, in general, the expression level of MHC-II genes was lower than that of MHC-I genes (**Extended Data Fig. 2a**). Notably, the expression of a proportion of*B2m*, *H2-K1*, and *Nlrc5*, was increased in EAE-OLGs already at early stages (**Fig. 2e and Extended Data Fig. 2a,b**). The chromatin accessibility of some of their corresponding genomic loci was accessible in OLG from CFA-Ctrl, and exhibited a further increase in the early stages of EAE (**Fig. 2e,f and Extended Data Fig. 2a**). In contrast, MHC-II genes had no or low expression in OLGs from CFA-Ctrl, and only showed increased expression and chromatin accessibility at the peak stage of EAE (**Fig. 2e and Extended Data Fig. 2a**). Importantly, MHC-I genes (such as *B2m*, *H2-D1*, and *H2-K1*) remained highly expressed at the late EAE stages, while the expression of most other MHC-I and -II genes were significantly downregulated compared to the peak stage. Nevertheless, chromatin accessibility at the promoter/gene body of both MHC-I and -II genes remained high at the late stage (**Fig. 2e,f and Extended Data Fig. 2a**). Thus, OLGs exhibit persistent chromatin accessibility of specific immune genes at late stages of EAE. Since inflammation is decreased at these stages, this immune chromatin accessibility persistence in OLGs might result from either epigenetic memory of peak immune-like states or from a mild inflammatory environment at late stages.

Our multiome data suggested that immune OLGs are present not only at peak EAE, but also at both early and late stages of EAE. We thus investigated with RNAscope *in situ* hybridization (ISH) targeting *Sox10* and *H2-Ab1* to understand where these imOLGs were located relative to EAE lesions. We defined lesions as white matter regions with a high number of cells due to inflammatory infiltrates. Consistent with the multiome results, the MHC-II^+^OLG (*Sox10*^+^*H2-Ab1*^+^) were observed in EAE spinal cord sections from the peak but also in both early and late stages, albeit in different proportions (**Fig. 2g,h**). The percentage of MHC-II^+^OLG was significantly higher at the peak stage (lesion: 5.33±0.23%, non-lesion: 5.32±0.23%) compared to both early stage (lesion: 0.49±0.849; non-lesion: 0.384±0.333%) and late stage (lesion: 2.54±0.23; non-lesion: 2.71±0.32%) in both lesion (P=0.0001) and non-lesion (P=0.0001) areas (**Fig. 2h**). However, there was no significant difference in the percentages of imOLG between lesion and non-lesion areas for all three stages (**Fig. 2h, P**<0.05). Altogether, MHC-I and -II genes maintain latent transcriptional and epigenetic states in OLG at late EAE stages, which may result in immune gene expression when facing recurrent inflammatory stimulus. While the expression of MHC-I in the early stages of the disease suggests a role in disease initiation, the expression of MHC-I and MHC-II in a subset of OLG at late stages might contribute to the chronic persistent disease state.

### Increase in the MOL2 population in peak and late EAE

MOLs have recently been shown to be heterogeneous, with specific populations exhibiting regional preferences and different susceptibility to spinal cord injury^5, 8, 12, 14–16^(. We identified MOL1, MOL2, and MOL5/6 populations based on their marker gene expression (**Fig. 3a and Extended Data Fig. 3a,b**), and the result of label transfer from our previous publication^5^ (**Extended Data Fig. 3c,d**). Based on distinct gene expression profiles, we could further subdivide the MOL1 into two (MOL1-α and β), MOL2 into twelve (MOL2-α, β, γ, δ, ε, ζ, η, θ, ι, κ, λ, and μ), and MOL5/6 into thirteen cell states/sub-populations (MOL5/6-α, β, γ, δ, ε, ζ, η, θ, ι, κ, λ, μ, and ν) (**Fig. 3a and Extended Data Fig. 4a**). These states present specific expression of gene modules, are involved in different biological processes (**Extended Data Fig. 4b**). For instance, when compared to other MOL2 populations, MOL2-ι, which mainly came from EAE at the early stage, expressed higher translation related genes, such as *Rpl6*, *Eef1b2*, and *Eif5a* (**Extended Data Fig. 4b**). Nervous system development genes, such as *Sema6a* and *Cntn4*, were enriched in MOL56-β and δ. A significant increase in the expression of translation related genes, such as *Rps27*, *Fau*, and *Uba52*, were observed in MOL5/6-γ compared to other MOL5/6 populations (**Extended Data Fig. 4b**). The majority of MOL2-α, β, γ, δ, and λ were derived from EAE late stage and majority of MOL2-ζ, η, θ, κ, and μ were derived from EAE peak stage (**Fig. 3b-d**). In addition, the majority of the cells in MOL5/6-β, ζ, η, and θ came from the CFA-Ctrl, while MOL5/6-γ and λ were mainly comprised of cells from EAE at the early stage, MOL5/6-ε and ν were mainly comprised of cells from EAE at the peak stage and cells from MOL5/6-κ and μ were drawn by EAE at the late stage (**Fig. 3c,d**). Overall, we observed a higher number of MOL2 at the peak and late stages compared to CFA-Ctrl and EAE early stage. This specific increase in the MOL2 population could arise from conversion of MOL5/6 into MOL2, MOL2 being more resilient than MOL5/6 to the neuroinflammatory environment, and/or preferential differentiation of OPCs into MOL2 in the context of EAE.

**Fig. 3.**
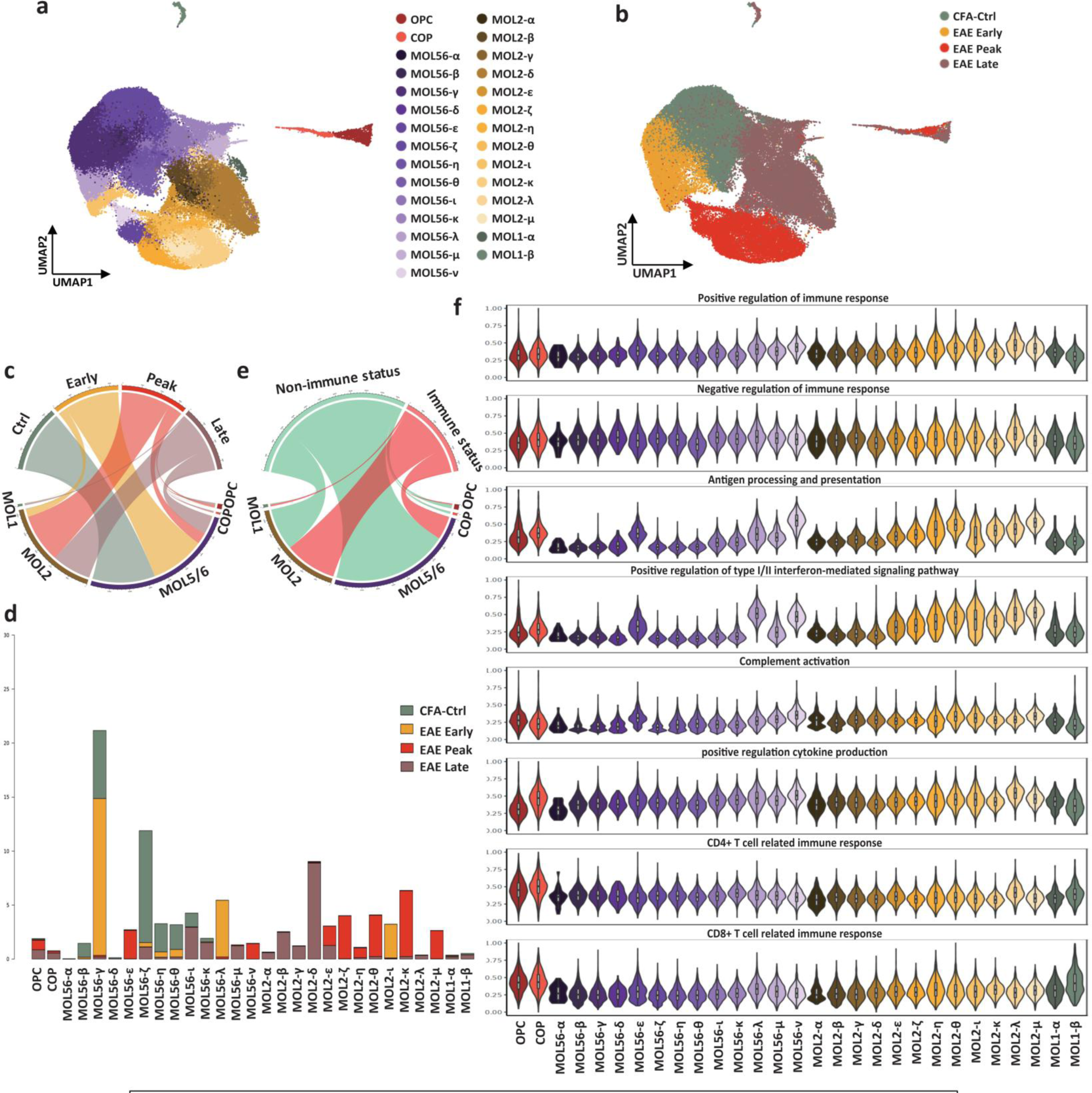
Stronger immune transcriptional responsiveness of the M0L2 mature oligodendrocyte population. a,b, Joint UMAP of OLG populations colored by sub-cell types (a) and time points (b). **c**, Circos plot showing the number of cells from different time points in OLG cell types, downsampled by time points (top levels). **d**, Bar plot of cell proportions in different conditions in each OLG sub-cell type. **e**, Number of cells with or without immune status for each OLG cell type. **f**, Scaled expression level of immune response related genes grouped by function in OLG sub-cell types.

### Stronger immune transcriptional responsiveness of MOL2

We then investigated whether any of these MOL populations were more prone to transition into immune-like-states in EAE. Indeed, 54.85% of MOL2 were identified as imOLG, while only 16.90% of MOL5/6 underwent this transition (**Fig. 3e**), indicating that MOL2 has higher immune responsiveness. Within the identified immune-related genes, some were more enriched in specific MOL populations, while others, involved for instance in cytokine and T cell immune response, showed similar expression levels among all MOL (**Fig. 3f**). We observed that genes related to positive regulation of immune response were enriched in MOL2-θ, ι, ν, and MOL5/6-ε, ν clusters (**Fig. 3f**), with the majority of these populations from EAE peak stage. Immune genes related to antigen processing/presentation and type I/II interferon-mediated signaling pathway were specifically enriched in MOL2-ε, ζ, η, θ, ι, κ, λ, μ and MOL5/6-ε, λ, μ, ν clusters. Genes related to complement activation had increased expression in MOL2-ν and MOL5/6-θ, μ, which again mainly came from EAE at the peak stage (**Fig. 3f**). Overall, our results suggest that MOL2 exhibit a stronger immune response to the neuroinflammatory environment than MOL5/6 and may play a more important role in the progress of the disease.

### Counter-balancing of immune responses in MOLs at early stages of EAE

Genes related to the negative regulation of the immune response were also expressed in nearly all MOL populations (**Fig. 3f**). However, a reduced expression was observed in MOL2-δ, ζ, κ, which mainly consisted of cells from the EAE peak and late stages (**Fig. 3f**). Moreover, we found that both the expression and chromatin accessibility at the promoter/gene body of *Cd274* (Programmed Death-Ligand 1, PD-L1) were increased at the early and peak stage in MOLs. PD-L1 is an immune checkpoint protein that interacts with the PD-1 receptor on immune cells and suppresses the immune response. The increase of PD-L1 expression may thus be a response to the exacerbation of the CNS inflammatory environment during the early and peak stages, stalling the autoimmune response in the disease. Interestingly, *Cd274* expression declined at the late stage even though its chromatin was still highly open (**Extended Data Fig. 5b,f**). This result suggests that a counter-balancing of negative immune regulation might occur at early stages of EAE in MOLs, with some sub-populations losing their ability to put a break on autoimmune response as the disease progresses to peak and late stages.

### Transitory regenerative responses in MOL5/6 during EAE

To explore whether other biological processes were dynamically modulated in OLG during the course of EAE, we performed differential gene expression across the different stages in different OLG cell types using a pseudobulk approach (see Methods). By comparing the gene expression in OPCs between CFA-Ctrl, EAE peak stage and late stage, the differential expression genes were divided into Type I (decreased at peak stage), Type II (increased at peak stage), and Type III (increased at late stage) (**Extended Data Fig. 5a**). As expected, genes related to immune response, such as *Nlrc5*, *Irgm1*, and *Iigp1* (Type II), were increased at the peak stage, followed by a significant reduction at the late stage (**Extended Data Fig. 5a**). However, many of these genes retained chromatin accessibility at their promoters and predicted enhancers, suggesting that they retain the epigenetic potential to initiate their transcription.

We also compared gene expression in MOL5/6 among the different stages and the differentially expressed genes of MOL5/6 were also divided into Type I (high expression in CFA-Ctrl), Type II (high expression at early stage), Type III (high expression at peak stage), Type IV (high expression at late stage) (**Fig. 4a**). The majority of the genes with high expression at the peak stage are immune-related, but their expression decreased significantly at the late stage (**Fig. 4a**). Strikingly, we found an increase in cholesterol biosynthetic process genes, such as Stearoyl-CoA desaturase-1 (*Scd1*) and Isopentenyl-Diphosphate Delta Isomerase 1 (*Idi1*), in MOL5/6 at the early stage (Type II) (**Fig. 4a,b**). Myelin is predominantly composed of lipids, and the formation of the myelin sheath necessitates highly coordinated levels of fatty acid and lipid synthesis process^17^. MOLs have been recently shown to be able to contribute in some extent to remyelination^18–21^. The elevated cholesterol biosynthetic-related genes suggest that there might be an adaptive increase of cholesterol biosynthesis at the initial stage of the disease to support the regeneration of myelin by mature oligodendrocytes, to compensate for EAE-driven demyelination. Interestingly, we also observe a slight and transient increase of specific myelin-related genes such as proteolipid protein (*Plp1)* and oligodendrocyte myelin glycoprotein (*Omg*) at this early stage (**Extended Data Fig. 5c**), albeit not other myelin genes. There was a subsequent decrease in the expression of these genes during the peak and late stages, as the disease progressed, suggesting a potential disruption of these putative remyelination processes. Nevertheless, we observed at late stages an increased expression of genes involved in RHO GTPase cycle in MOL5/6, such as *Rgs16* and *Arhgef19* (**Fig. 4a**), involved in actin cytoskeleton and microtubule processes, which play a role in myelination^22^, further suggesting that MOL5/6 might activate gene regulatory programs associated with regeneration during the early stages of EAE.

**Fig. 4.**
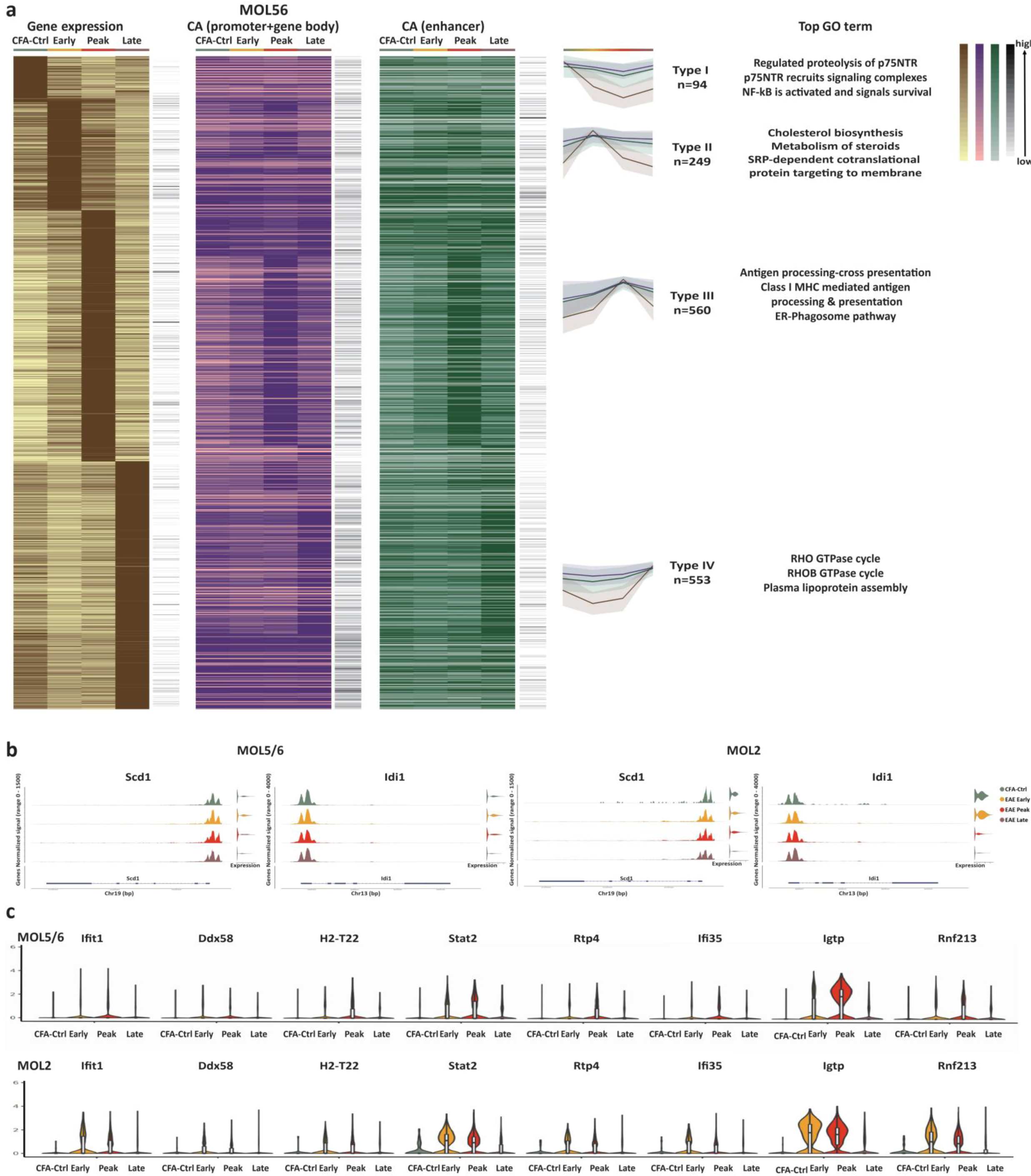
Increase of cholesterol biosynthetic processes in MOL5/6 and immune processes in M0L2 at early stages of EAE. **a**, Heatmaps of differentially expressed genes (in brown) between different time points in MOL5/6, the chromatin accessibility at promoter and gene body (in purple) and chromatin accessibility at enhancer regions (in green) of the same gene (left), with the top GO terms of each group of genes (middle). The black column on the side represents the gene raw counts. The line plots represent the mean of the normalized and scaled gene expression (line in brown), the chromatin accessibility at promoter and gene body (line in purple), and chromatin accessibility at enhancer regions (line in green) of genes in different groups (right). **b**, Normalized chromatin accessibility (left) and Iog2 expression (right), of genes related to cholesterol biosynthetic process in MOL5/6 (left) and M0L2 (right). **c**, The expression of representative immune related genes (belong to M0L2 Type II differentially expressed genes) in MOL5/6 (top) and M0L2 (bottom).

### MOL2 develops immune transcriptional responses to EAE earlier than MOL5/6

We could also identify four groups of genes with differential expression between different time points for MOL2 (**Extended Data Fig. 5d**). The top GO terms associated with Type I genes (high expression in CFA-Ctrl) included ‘axon guidance’ and ‘nervous system development’, with genes like *Epha4*, *Sema4a*, and *Tenm2*. The decreased expression of these homeostatic CNS-related genes at the early and peak stages could be consistent with the observed aggravating symptoms of EAE mice during the disease (**Extended Data Fig. 5d**). In Type II genes with high expression at the EAE early stage, as in MOL5/6, we found that cholesterol biosynthetic process related genes, such as *Scd1* and *Idl1*, also had a transitory increase in MOL2 at EAE early stage compared to CFA-Ctrl (**Fig. 4b and Extended Data Fig. 5d**). The initial upregulation of cholesterol-related genes was followed by a significant decrease during the peak stage of EAE and with limited or no recovery at the late stage (**Fig. 4b**), suggesting that transitory regenerative response in early stages occurs in the different subtypes of mature oligodendrocytes.

Genes involved in translation and RNA processing related genes, such as *Rpl29*, *Rps17,* and *Eif1a*, also displayed increased expression in MOL2 during the early stage of EAE, followed by a decrease at the later stages (**Extended Data Fig. 5d,e**), suggesting a highly active metabolic state of MOL2 at the early stage of EAE. In contrast to MOL5/6, we found that a subset of immune genes (such as *H2-T22*, *Stat2*, and *Rtp4*) had increased in MOL2 at the early stage (**Fig. 4c**). This finding further suggests that MOL2 may play a more important role in the disease progression at the initial stage. Many genes with high expression in MOL2 at the peak stage were immune-related genes (Type III, such as *Tap1*, *H2-T23*, and *Itga9*), which is consistent with the exacerbated immune response and higher number of imOLG at the peak stage. (**Extended Data Fig. 5d**). Interestingly, we also found genes related to developmental and axon guidance, different from Type I genes, exhibited high expression in MOL2 at the late stage (Type IV), such as *Smarca2*, *Map1b*, and *Gas7* (**Extended Data Fig. 5d**). Thus, the environment at the late stages of EAE, when inflammation resolves, might trigger the induction of neural developmental gene programs in MOL2. In summary, our multiome data indicates that the MOL2 initiates, as MOL5/6, a cholesterol biosynthesis program in response to the arising neuroinflammatory environment, but transitions to an immune-like state earlier than MOL5/6 during the course of EAE.

### Early increase in chromatin accessibility at promoter/gene bodies of immune genes in MOL2 in EAE

Differences between MOL2 and MOL5/6 at the transcriptional level might correlate with changes at the epigenetic level. We thus compared the list of genes with differential chromatin accessibility at promoter/gene body regions and genes with differential expression across different stages and found that 188 genes in MOL2 and 115 genes in MOL5/6 showed significant changes in both expression and chromatin accessibility **(Fig. 5a)**. For both MOL2 and MOL5/6, the function of the majority of these genes were immune-related, with top GO terms of ‘‘antigen processing-cross presentation’’ and ‘‘ER-phagosome pathway’’ **(Fig 5b, Extended Data Fig. 6e,f)**. This indicated that immune-related genes tend to have significant changes in both gene expression and chromatin accessibility.

**Fig. 5.**
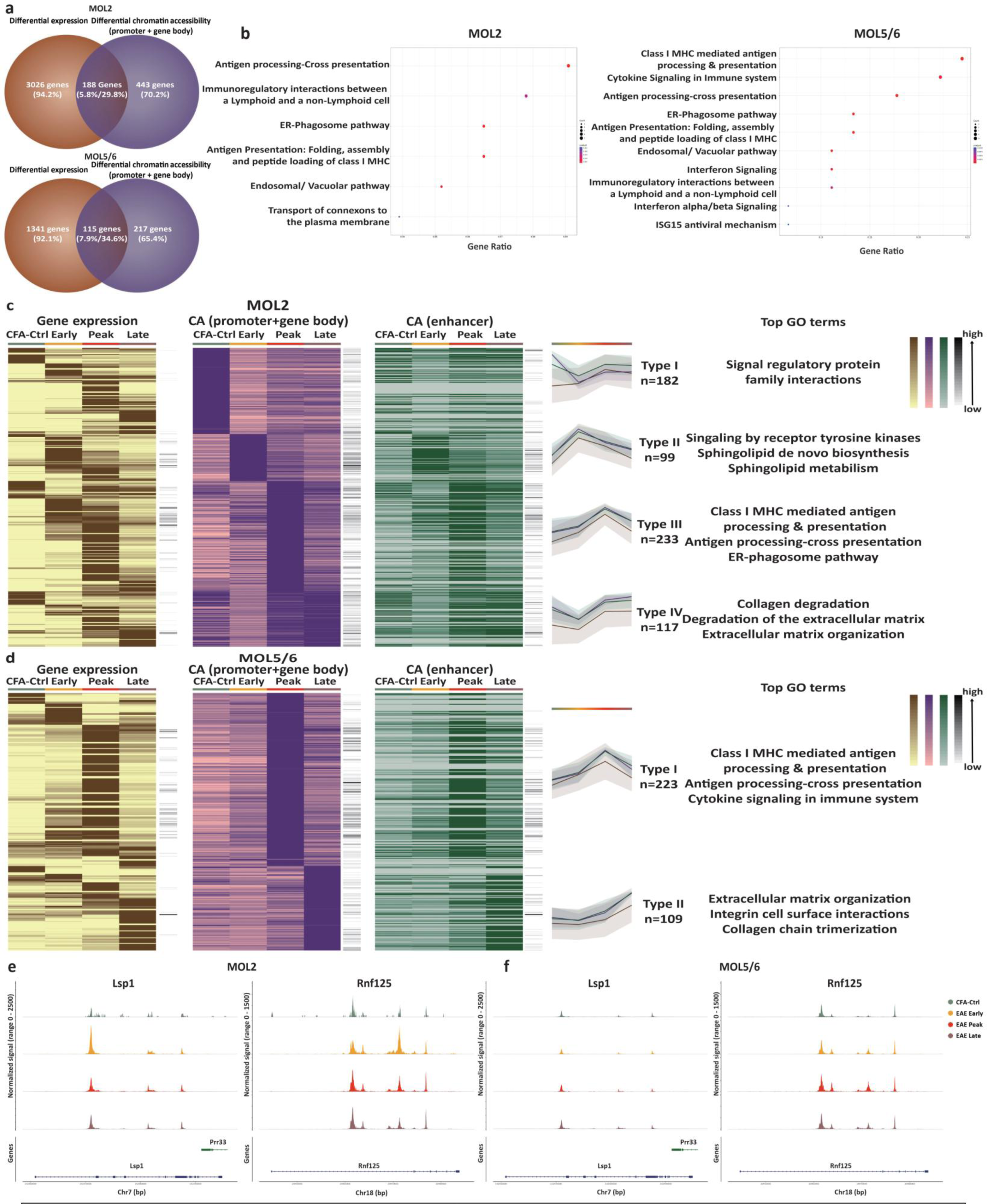
Earlier and stronger epigenetic immune response in M0L2 than MOL5/6. **a**, Venn diagram showing the number of genes with differential expression and/or differential chromatin accessibility among different disease stages in MOL2 (top) and MOL5/6 (bottom). **b**, Top gene ontology by biological process terms for genes with both differential expression and differential chromatin accessibility in MOL2 and MOL5/6 among different disease stages. **c, d**, Heatmaps of differential chromatin accessibility at promoter and gene body (in purple) between different time points in MOL2 (c) and MOL5/6 (d), of gene expression (in brown) and chromatin accessibility at enhancer regions (in green) of the same gene (left), and the top GO terms of each group of genes (middle). The black column on the side represents the gene raw counts. The line plots represent the mean of gene expression (line in brown), the chromatin accessibility at promoter and gene body (line in purple), and chromatin accessibility at enhancer regions (line in green) of genes in different groups (right). **e**,f, Normalized chromatin accessibility of Lspl and Rnfl25 in M0L2 (c) and MOL5/6 (d) at each time point.

We then explored the dynamics of chromatin accessibility both at promoter/gene body and at distant enhancer regulatory regions, during the EAE disease course. Gene loci displaying differential chromatin accessibility at the promoter/gene body regions between different time points in MOL2 could be divided into 4 types: high chromatin accessibility in CFA-CTRL compared to any EAE stage (Type I), high chromatin accessibility at the early stage (Type II), peak stage (Type III), and late stage (Type IV) **(Fig. 5c)**. In contrast, only two different types of chromatin accessibility variation patterns were found in MOL5/6: high chromatin accessibility at the peak stage (Type I) and at the late stage (Type II) **(Fig. 5d)**.

As observed at the transcriptional level, we found that the chromatin accessibility at promoter/gene body regions of genes involved in myelination-related processes, as sphingolipid de novo biosynthesis and metabolism^23^, at the early stage in MOL2 (Type II, **Fig. 5c**). We did not observe differences in such pathways in MOL5/6, which might be related to an already increased chromatin accessibility at promoter/gene body regions in CTRL-CFA conditions (**Fig. 5d**). Some immune system process related genes, such as *Lsp1* and *Rnf125,* also presented increased chromatin accessibility in MOL2 at the early stages **(Fig. 5e)**. However, we did not find the increase of chromatin accessibility of these immune genes in MOL5/6 at the early stage (**Fig. 5f**). Nevertheless, we found that there was a group of antigen processing/presentation with high chromatin accessibility at the peak stage in both MOL2 and MOL5/6, such as *Irgm1*, *Igtp*, and *Gbp7* **(Extended Data Fig. 6a,b)**. Some immune system process-related genes, such as *C3*, *Lbp*, *Isg20*, only showed significantly increased chromatin accessibility at the peak stage in MOL2 but not in MOL5/6 **(Extended Data Fig. 7c,d)**, further indicating that MOL2 triggers a more robust opening of the chromatin at immune genes upon EAE than MOL5/6.

### Enhancer-mediated regulation of immune genes in MOL5/6 and MOL2 in EAE

In addition to promoters, enhancer regions distal to transcription start sites have been shown to regulate the transcription of associated genes. Recently, super-enhancers, clusters with high levels of transcription factor binding, and domains of regulatory chromatin (**DORCs**), regions with high density of ATAC-Seq peaks, have been suggested to have key roles in modulating gene expression^24^. To explore the role of DORCs in the course of the disease in EAE, we were able to define 162 differential accessible domain-regulated genes in MOL2 and 54 differential accessible domain-regulated genes in MOL5/6 with an exceptionally large (≥5) number of significant peak-gene associations as DORCs (**Fig. 6a and Extended Data Fig. 7a**).

**Fig. 6.**
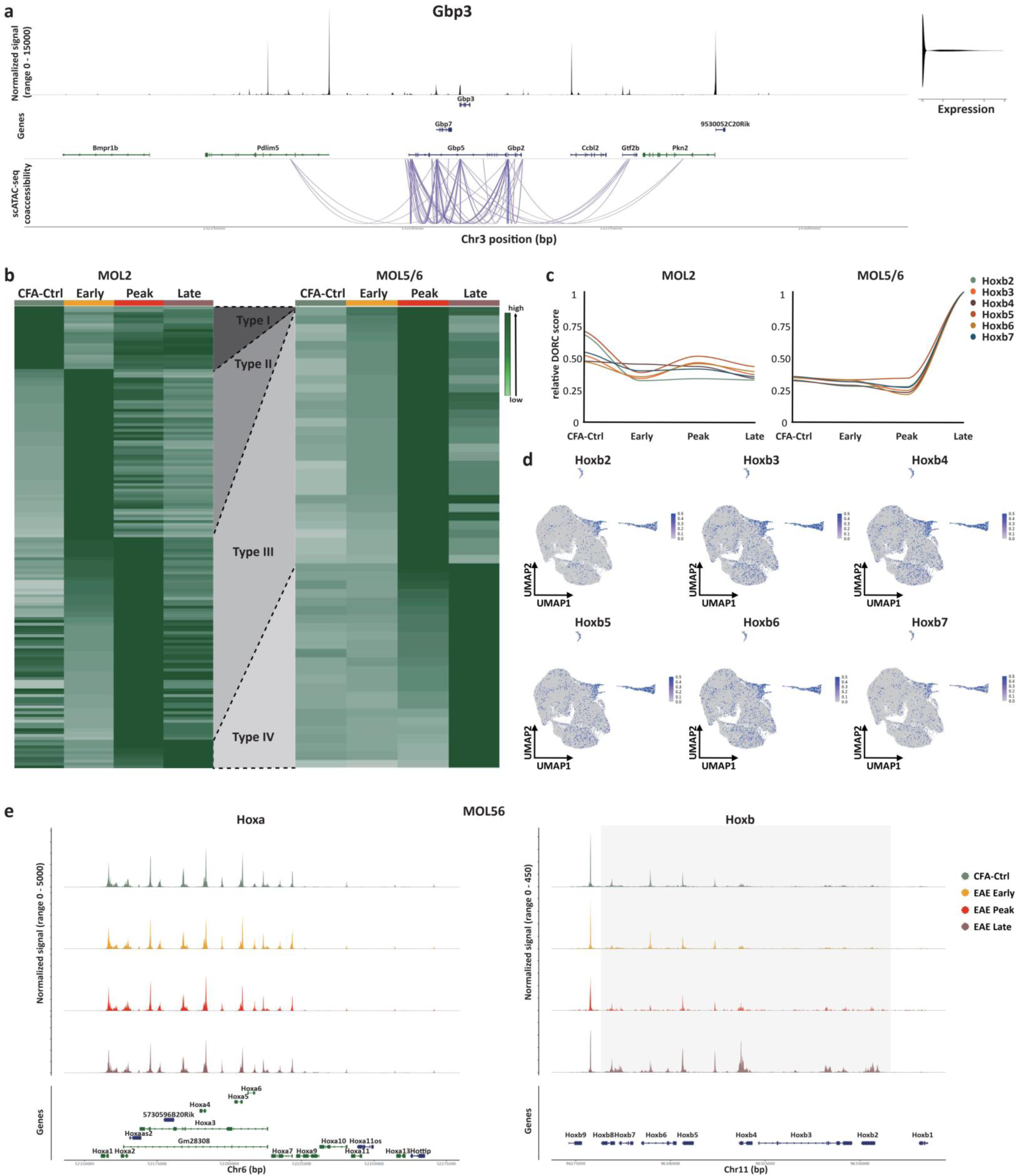
Increased chromatin accessibility at Hoxb family DORCs in MOL5/6 at the late stage of EAE. **a**, Representative DORC (Gbp3). The genomic track represents the OLG bulk accessibility of Gbp3 while the links denote the significant correlation (pvalue < 0.05) between peaks and Gbp3 (± 500 kb from TSSs). The violin plot shows Gbp3 Iog2 expression in OLG. **b**, Heatmaps of normalized and scaled DORC score of 216 genes in MOL2 and MOL5/6 at different stages, DORCs were classified into three types: type I with decreased activity at peak and late stages, type II with increased activity at the peak stage and type III with increased activity at the late stage. **c**, The normalized and scaled DORC score of representative cell development related type IV DORCs in MOL2 and MOL5/6. **d**, Feature plots showing DORC scores of Hoxb2, Hoxb3, Hoxb4, Hoxb5, Hoxb6, and Hoxb7. UMAP coordinates and population distribution as in Figure 3a. **e**, Normalized chromatin accessibility of Hoxa (left) and Hoxb (right) genes in MOL5/6, grey box highlight Hoxb genes in d.

We found four types of DORCs in MOL2, among which 22 domains were classified as Type I, which had high activity in CFA-Ctrl and early stage but decreased at peak and late stages, and 59 domains were classified as Type II, which showed high activity at the early stage (**Fig. 6b**). Similar to the chromatin accessibility at the gene body and promoter region, we also found some immune-related genes such as *Ciita*, *Ido1*, and *Irf7* showed increased enhancer activity at early stage in MOL2 (Type II), which was also consistent with the early expression of immune gene MOL2 (**Fig. 6b and Extended Data Fig. 7b)**. In contrast, we only found two types of DORCs in MOL5/6, type III (high activity at peak stage) and type IV (high activity at late stage), consistent with the chromatin accessibility at the promoter/gene body level (**Fig. 5a,b**). Nevertheless, Type III DORCs in both MOL2 and MOL5/6 were associated with genes involved in immune processes, with *Gbp3*, *Iigp1*, and *Irgm1* included (**Fig. 6b and Extended Data Fig. 7c**).

### Increased chromatin accessibility at *Hoxb* regulatory regions in MOL5/6 at the late stage EAE

Interestingly, Type IV DORCs showed gradually increased activity at the late stages of the EAE disease course (**Fig. 6b**). In particular, we found increased DORCs regulating a group of *Hoxb* genes, such as *Hoxb2*, *Hoxb3* and *Hoxb4,* in MOL5/6 but not MOL2, (**Fig. 6c-e and Extended Data Fig. 7d,e**). The increased activity was only found in *Hoxb* but no other Hox family gene regions (**Fig. 6e and Extended Data Fig. 7e**) Hoxb2 was demonstrated to be essential for oligodendrocyte patterning^25^. Thus, the increased chromatin accessibility of these type IV developmental-related DORCs in MOL5/6 suggests that MOL5/6 might have primed transcriptional programes compatible with nervous system repair and remyelination and promote during the late stages of the disease in EAE.

### MultiVelo analysis indicates divergence responses of MOL2 and MOL5/6 to the evolving disease environment in EAE

Single-cell RNA velocity is a powerful tool for understanding gene regulation and dynamics of cellular processes through various states by computationally estimating the rate of change in spliced and unspliced transcripts over time in individual cells^26^. While RNA velocity leverages splicing and RNA turnover to infer cellular transitional dynamics, another key component for these dynamics are changes in the epigenomic landscape during cell transitions. To explore these coupled dynamics of OLG during the disease course, we applied MultiVelo^27^, a tool that integrates transcriptomics and epigenomics datasets to estimate cell-fate predictions. This tool is anchored in two models for the correlation of gene expression and chromatin accessibility changes within the latent timeline (**Fig. 7a and Extended Data Fig. 8a,b**). In these models, it is possible to define, for a given gene, the state of each cell into one of the following four phases, according to the levels of chromatin accessibility, unspliced pre-mRNA and spliced mature mRNA: priming (brown, chromatin is opening but transcription is not initiated), coupled-on (pink, chromatin is opened and transcription is initiated), decoupling (dark blue, M1: chromatin accessibility starts closing before the end of the transcription, M2: chromatin is opened but transcription repression begins), and coupled-off (light blue, chromatin is closed and the number of unspliced reads is dropping)^27^.

**Fig. 7.**
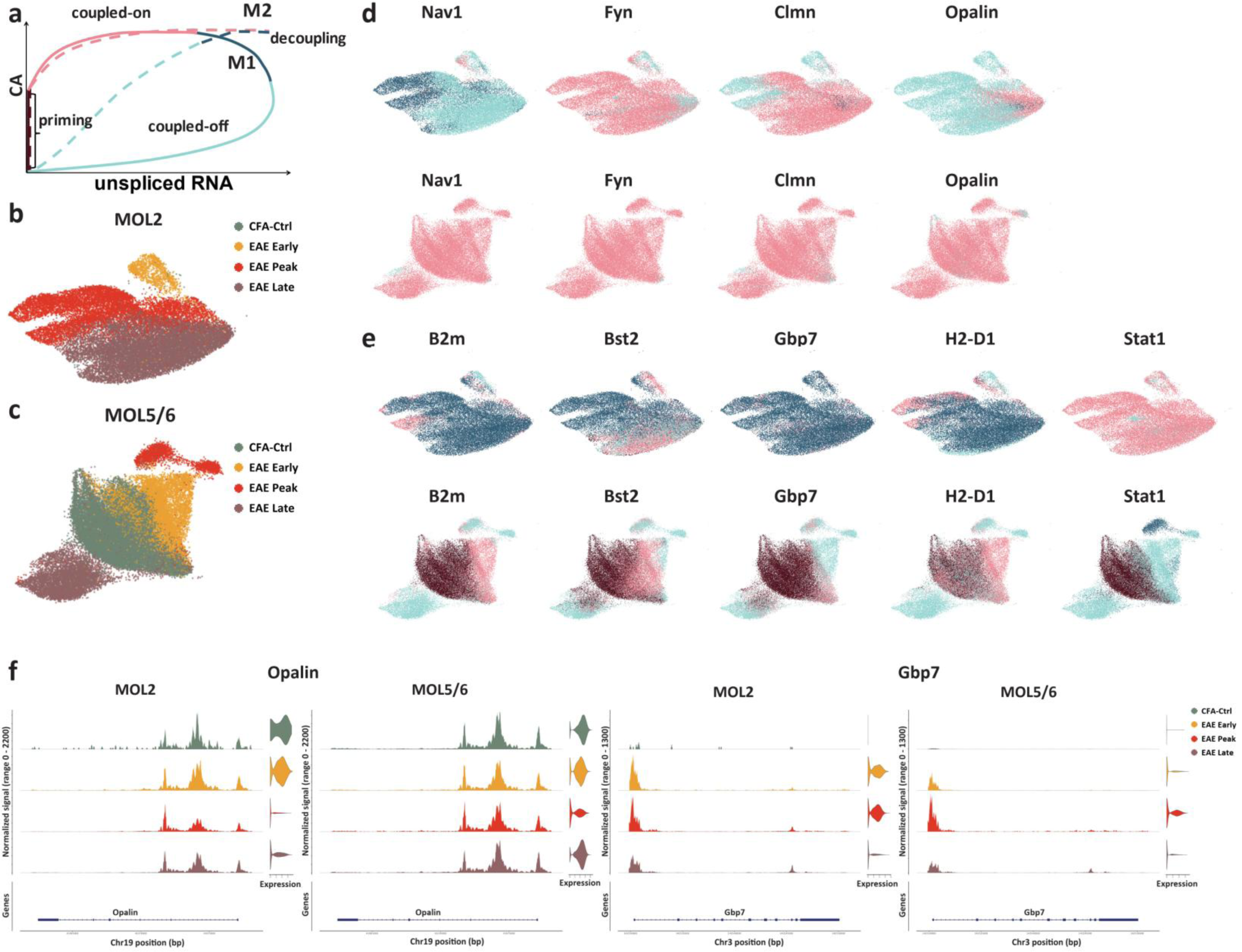
MultiVelo analysis indicates divergence responses of MOL2 and MOL5/6 to the evolving disease environment in EAE **a**, Gene phase portraits predicted by MultiVelo for model 1 (Ml, full line) and model 2 (M2, dotted line) genes. **b, c**, MultiVelo UMAPs of MOL2 (b) and MOL5/6 (c) colored by time points. **d,e**, UMAP of MOL2 and MOL5/6 colored by gene state assigned by MultiVelo for representative neuron system development (d) and immune response related genes (e). Navi, Fyn, Clmn, Opalin, and Statl are classified as Ml genes, B2m, Bst2, Gbp7, and H2-D1 were classified as M2 genes, f, Normalized chromatin accessibility (left) and Iog2 expression (right) of Opalin and Gbp7 in MOL2 and MOL5/6.

We found that canonical OLG genes such as *Opalin*, *Fyn*, and *Nav1*, showed coupled-on phases in MOL5/6 in both CFA-Ctrl and in EAE from early to late stages (**Fig. 7c-e**). However, in MOL2, *Opalin* exhibited a coupled-off phase, in particular at early and peak EAE stages. MOL2 cells present a coupled-on phase only at late stages (**Fig. 7b,d,e**). *Opalin* (Tmem10) has been shown to play a role in oligodendrocyte differentiation^28^. Thus, similar to our previous results, Multivelo analysis suggests a differential regenerative response of MOL2 and MOL5/6 in the context of EAE.

Immune-related genes, such as *B2m*, *Bst2, H2-D1, Stat1* and *Gbp7*, were observed to transition between the coupled-on and decoupling phases in the majority of MOL2 cells, which indicated a highly open chromatin level and transcripts (**Fig. 7b,d**). However, these immune genes showed a chromatin priming phase in MOL5/6 from CFA-Ctrl, followed by a transient coupled-on phase at EAE early stage, but transitioned into coupled-off at the peak and late stages, indicating a significant reduction in chromatin accessibility and gene expression of these immune genes at the chronic stage of the disease (**Fig. 7b**). Transition to an immune -like state has been previously shown to be incompatible with differentiation in OPCs^6^, and our data suggest that in mature oligodendrocytes in EAE the enhanced immune-like state of MOL2 is distinct to a putative more remyelination-prone state of MOL5/6. Thus, these results further indicate a divergence in the response of different MOL populations to the evolving disease environment in EAE.

### Gene regulatory network analysis indicates sharp changes in transcription factor activity in MOLs during EAE disease progression

To obtain insights into the molecular mechanisms mediating the divergent MOL responses in the different stages of EAE, we inferred the global gene regulatory network (**GRN**)^29^ for MOL5/6 and MOL2. We first selected differentially accessible regions specific for each time point and MOL (see Methods). Then, the regulatory regions were scanned for transcription factor (**TF**) binding sites to fit the Generalized Linear Model (GLM) between candidate regions with TF motifs and the expression of the possible target genes. Based on the model, we selected TFs that show expression and ranked them by their higher activity at each timepoint (see Methods). We observed changes in the activity of the predicted TF between the different stages of EAE (**Fig. 8a-c and Extended Data Fig. 9**). For instance, TFs associated with immunoregulatory functions, such as *Stat1* and *Irf7,* presented low activity in both MOL2 and MOL5/6 in CTRL-CFA, dramatically increase activity at the early stages and reached top positive activity at the peak stage, consistent with our prior study^12^. Interestingly, while *Irf7* TF activity returned to basal levels at the late stages, *Stat1* TF activity remained at a relatively high level (**Fig. 8a-c and Extended Data Fig. 9**).

**Fig. 8.**
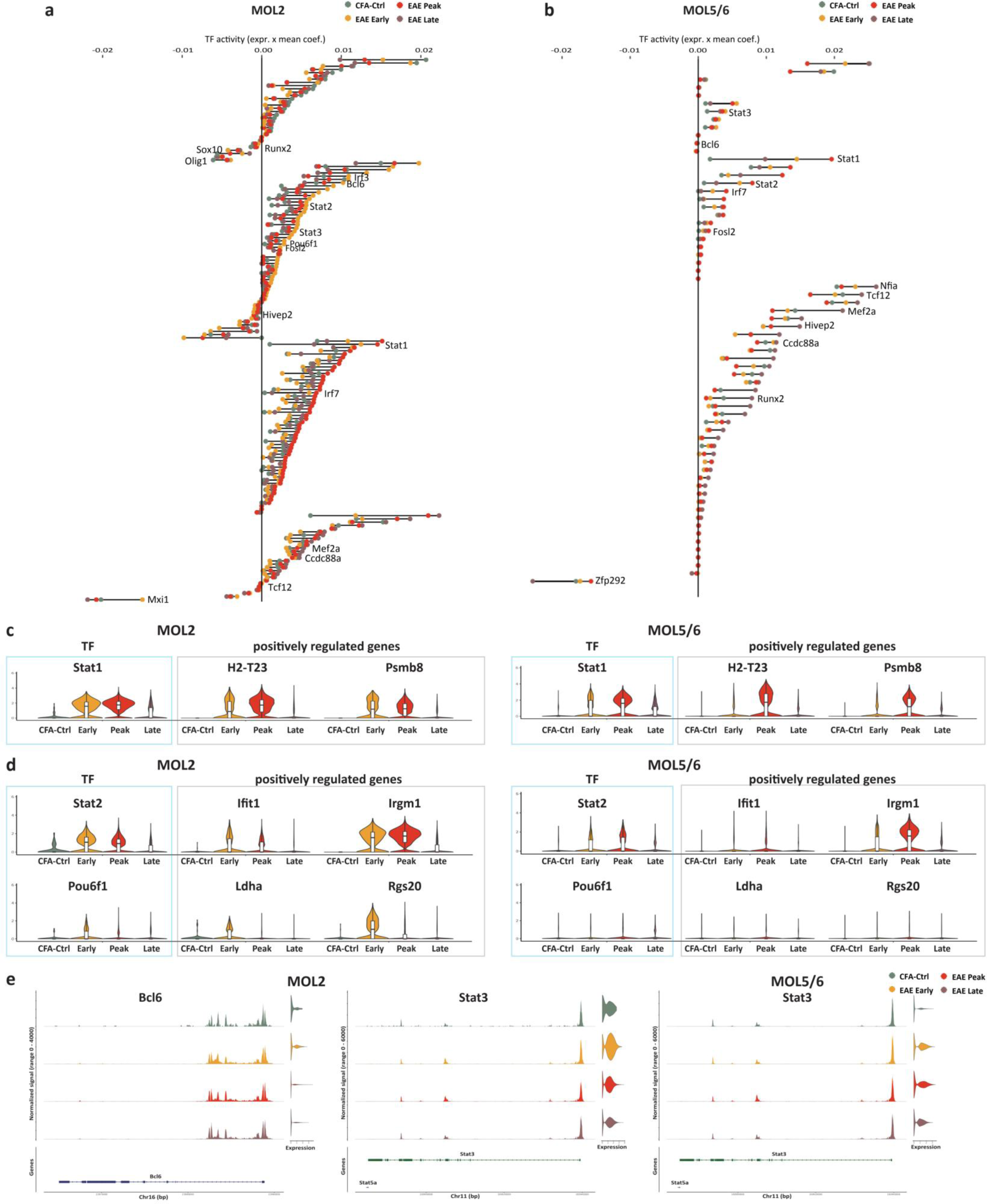
Transcription factor activity foresees the immunosuppression potential of MOL **a,b**, Rankend TF activities of the predicted TFs at different stages in M0L2 (a) and MOL5/6 (b). x axis: TF activity represented with a dot colored by stage and connected with a line. TF activity was calculated as the mean coefficient multiplied by the average expression in each time point., y axis: transcription factors ranked based on the TF activity in each time point, (zoom in view in supplementary figure. 9). **c**. Violin plots showing the expression of Statl TF and a selection predicted target genes that are positively regulated by Statl at different stages in MOL2 (left) and MOL5/6 based on the inferred model (right). **d**. Violin plots showing the expression of Stat2, Pou6fl and their predicted positively regulated genes (Stat2: Ifitl and Irgml; Pou6fl: Ldha and Rgs20) at different stages in M0L2 (left) and MOL5/6 (right). **e**. Normalized chromatin accessibility (left) and Iog2 expression (right) of Bcl6 and Stat3.

We found a higher number of TFs, acting mainly with positive activities, at the late stage in MOL5/6 (38 TFs) compared to MOL2 (27 TFs). Among these TFs, *Ccdc88a* and *Mef2a* (**Fig. 8a,b**), which are related to nervous system development, showed higher activity at the late stage in both MOL2 and MOL5/6. However, within MOL5/6 predicted TFs, we found an increased number of candidates related to nervous system development mainly acting in a positive manner at the late stage, compared to the CFA-Ctrl and the other stages. These TFs include *Nfia* and *Tcf12*, which are important TFs involved in OL differentiation and maturation^30, 31^ (**Fig. 8b**). Some cell differentiation-related TFs, such as *Runx2* and *Hivep2*, were also found mainly acting at late stage compared to other stages in MOL5/6 (**Fig. 8b**). These results further underscore the stronger regenerative potential in MOL5/6 during the late stages of the disease.

### Transcription factor activity is consistent with the immunosuppression in MOLs in early stages of EAE

Most TFs were actively regulating in EAE compared to CFA-Ctrl in MOL2 and MOL5/6 (**Fig. 8a-c and Extended Data Fig. 9**). A subset of TFs associated with the positive regulation of immune responses, such as *Stat2*, *Fosl2*, *Pou6f1*, and *Irf3*, presented their highest activity at the EAE early stage in MOL2 (**Fig. 8a,d**). This result is consistent with the amplified and earlier immune transcriptional responsiveness observed in MOL2. Interestingly, we also found that transcription factors such as *Bcl6* and *Stat3*, which have immunosuppression potential^32,33^ showed higher positive activity in MOL2 at the early stage of EAE, compared to the peak and late stages (**Fig. 8a,e**). Notably, *Stat3* was acting in the same manner in MOL5/6 (**Fig. 8b,e**). Taken together with the increased expression of PD-L1 in MOL from EAE, this result suggests that MOLs may act as regulators in mitigating autoimmune responses, particularly at the early disease stages.

## Discussion

The immune plasticity of OLG was first reported about 40 years ago^34^. In addition, MHC-I and -II expression in OLG under inflammatory conditions was further observed in many subsequent studies^35–38^. Recently, with the emergence of single-cell transcriptomics, the concept of immune OLGs has been confirmed and further expanded to neuroinflammation in the context of MS; Alzheimer’s disease or aging in several studies^6, 7, 39, 40^. We have recently shown that MHC and other immune-associated genes are highly expressed in EAE at the peak stage, with primed chromatin accessibility of some of these genes already in CFA-Ctrl^5, 12^. In the current study, we applied multi-omics scRNA-seq and scATAC-seq, which allowed us to obtain a comprehensive view of the cellular states of OLG during the disease process in EAE, including the onset and the chronic disability stages of the disease.

Our results show that several immune-related genes and their chromatin accessibility levels are already significantly elevated in OLG at the early stage of EAE, when symptoms start emerging. Furthermore, we observed that the chromatin loci of immune-related genes remain highly accessible in the late stage of the disease. The induction of OLG with immune profiles occurs at the early stages of EAE, when lesions begin to form but not yet fully developed or not prevalent^41^. We also observe that the percentage of MHC-II^+^OLGs is similar inside and outside lesions, indicating that the lesion environment is not essential for the transition of OLGs to immune-like states. Recent spatio-temporal mapping of cells in EAE suggests that the induction of disease-associated glia precedes lesion formation and that the disease-associated MOLs are in close proximity to the disease-associated microglia and astrocytes^41^. Thus, it is plausible that the induction of immune OLGs might be mediated not only by infiltrating immune cells, but also by other disease-associated glia or other inflammatory environmental factors.

The presence of immune OLGs at early, peak and late stages of EAE could suggest differential functions at these distinct timepoints. We have previously shown that immune OLGs express both activating and inhibitory immune checkpoint molecules^12^, and our results are consistent with the activation of immune repressive pathways in MOLs at the early and peak stages of the disease. Thus, OLGs could in principle activate or, alternatively, block immune responses in the context of EAE. At the early stages, immune OLGs might act by modulating (preventing or initiating/amplifying) the initial neuroinflammatory events underlying the aetiology of the disease. At the peak stages, given the presence of high numbers of professional antigen presenting cells such as microglia and macrophages, the role of immune OLGs might be more subtle and modulatory of the function of these other immune cells. We observe the persistence of MHC-I and MHC-II in OLGs in the late stages of EAE, when immune OLGs might be involved in the persistence of the disease, although a role in reducing inflammatory responses is also possible.

In the most common course in MS, RRMS patients suffer from multiple remissions and relapses in the disease course, and the symptoms can end up worse than before with each relapse^1^. Therefore, the observed chromatin accessibility memory of immune-related genes in OLG may contribute to faster and stronger immune gene expression upon the next wave of stimulation during the relapse stage in MS patients, which can result in the progressive enhancement of myelin-specific immune responses and demyelinating. These findings suggest that OLGs play an active role in the immune response in EAE, and the early initiation and persistence of the immune response chromatin memory in OLGs may contribute to the chronicity of the disease and the difficulty in treating demyelinating diseases.

The heterogeneity of the OL lineage in development has been reported in our previous study^14^, and several subsequent studies have further confirmed the transcriptional and spatial preferences of distinct subpopulations of OL in development and disease^5, 8, 15, 16^. In our analysis, we identified previously reported mature resting subtypes, specifically MOL1, MOL2, and MOL5/6. The percentage of cells with an immune profile was higher in MOL2 than in MOL5/6. The expression of immune-related genes in MOL5/6 decreased significantly at the late stage compared with the early stage and peak stage. Taken together, the immune characteristics of MOL2 in the spinal cord of EAE mice are stronger than that of MOL5/6. MOL5/6 are more enriched in the gray matter of the spinal cord, while MOL2 are more preferentially reside within the white matter of the spinal cord, where the immune infiltrates and lesions are the most prominent^15, 42^. The difference in the domain distribution between MOL2 and MOL5/6, in combination with intrinsic factors, may thus contribute to the heterogeneity of MOL in response to the inflammatory environment and higher immune characteristics of MOL2.

Remyelination is a spontaneous regenerative process in which the CNS strives to repair itself. Several factors, including cell senescence, depletion of the OPC pool, and environment signaling changes, have been proposed as potential elements contributing to the eventual failure of the myelin repair mechanisms in MS^43^. Many studies have demonstrated that remyelination also occurs within MS lesions while most of the time it is insufficient^44–46^, and it is still unclear which cells are responsible for remyelination and at what stage. OPCs are viewed as an origin of OLs and have been identified within lesions in post-mortem CNS tissues from MS patients^47^. However, it remains unclear whether OPCs in lesions develop into fully differentiated OLs capable of remyelination, and to what extent newly generated and existing OLs contribute to myelin regeneration. The remyelination process usually involves the sequential activation and recruitment of OPCs to demyelination sites followed by their differentiation into myelinating oligodendrocytes. Studies with mouse demyelination models have demonstrated successful remyelination attributed to new OLs generated from OPC proliferation and differentiation^20, 48, 49^. However, some studies have also shown that survival MOLs in the demyelinating region also contributes to the remyelination process^19, 21, 50^. Recent C^14^ birth dating in human MS patients^18^and live imaging in mouse^51^and zebrafish^20^, suggest that remyelination in the context of demyelination can be promoted by MOLs, albeit in a non-efficient manner.

Our results revealed a transitory increase of the cholesterol biosynthesis process in MOLs at the early stage of EAE, in particular in MOL5/6. Consistent with the cholesterol biosynthetic process genes, some genes that are closely associated with OL myelination, such as *Plp1*, and *Omg*, exhibited a slight and short-term increase of expression at the early stage, but followed by a decrease at the peak stage and without significant recovery at the late stage (**Extended Data Fig. 6b**). These findings suggest that existing MOLs could respond rapidly during a very early demyelination stage and increase their cholesterol synthesis to compensate for demyelination caused by the myelin specific immune response. These cholesterol biosynthesis genes decrease as the disease progresses, which may be due to the diminished remyelination ability of the MOLs or cell death after multiple bouts of demyelination. MultiVelo analysis also indicated a coupled-on transcriptional and epigenetic state for OL differentiation genes in MOL5/6 at all stages of EAE. Therefore, promoting myelin regeneration in the existing MOLs, in particular MOL5/6, at the early stage and in the surviving MOL at the late stage, may be a meaningful therapeutic target of MS.

We observed notable disparities in CNS development functions between MOL2 and MOL5/6. We saw that chromatin accessibility of some genes related to CNS development always stay in a coupled-on stage in MOL5/6 but not in MOL2, which suggests a more significant role of MOL5/6 in myelin maintenance and stability than MOL2 in the disease, especially at the chronic stage. However, our results do not provide a definitive answer regarding whether MOL5/6 at the late stages are newly generated or resilient cells that were already present before disease development. One hypothesis is that MOL5/6 are newly generated and possess a stronger ability for CNS development. Alternatively, it is still possible that these MOL5/6 survive in the severe inflammatory environment, and contribute to remyelination once the inflammation in the CNS is diminishes. To address this question, lineage tracing experiments need to be conducted in future studies.

Some studies reported that immune responses are different in many aspects between males and females in both MS and its animal models^52, 53^. Since most of our samples were mixed with one male mouse and one female mouse, we created a sex prediction model based on the expression of sex-related genes (see methods). This sex prediction model was validated with samples containing cells from only male or female EAE mice, with an accuracy of 95.3% on the validation dataset. There was better prediction accuracy towards the male sample (99.48%) than in the female sample (86.13%) (**Extended Data Fig. 10a,b**). We applied this sex prediction model to the entire dataset, annotated the sex of the cells (**Extended Data Fig. 10c,d**), and compared the differences between males and females OLGs based on the sex prediction results. For OLG sub-populations, apart from sex related genes, such as *Xist, Tsix, Uty, Eif2s3y, Ddx3y,* and *Kdm5d*, no other genes were found differentially expressed between male and female (**Extended Data Fig. 10e**). Accordingly, we found no major differences between the OLGs of males and females at the gene expression level in our data. We also did not observe a significant difference in the percentages of imOLGs between male and female mice (**Extended Data Fig. 10f,g**). Thus, our data indicates that the OLG response to the neuroinflammatory environment in EAE is not characterized by sexual dimorphism.

Our study provides a unique single-cell multiome resource for understanding the intricate mechanisms underlying the gene regulation and chromatin accessibility of OLGs in the context of MS using an animal model. In particular, it sheds light on the distinct fates of MOL2 and MOL5/6 during disease evolution, indicating that therapeutic strategies targeting MOLs need to be specific for the different populations. Moreover, our analysis framework serves as a blueprint for making discoveries using single-cell multi-omic data. The road to fully understand the complex interplay of transcriptomics and epigenomics in MS requires further exploration, such as spatial resolution, which may provide a more comprehensive and accurate picture of the disease.

### Limitations of the study

Our dataset gives a unique view of the epigenomic and transcriptomic events underlying the response of MOLs to the evolving inflammatory environment that occurs in MS. Nevertheless, EAE is an outside-in model of MS that relies mainly on the recruitment of myelin-specific T-cells. While mimicking many aspects of MS, other factors might be operational at the disease niche in MS, and thus further influence MOL response. The use of alternative MS mouse models, such as transgenic spontaneous model and virus-induced/toxic-induced demyelination model, will elucidate whether this is indeed the case.

Our multiome analysis gives insights into the epigenetic potential and memory in MOLs during the time course of the disease, at the onset of the disease, when symptoms are starting to manifest, at the symptomatic peak and when the disease stabilizes at the chronic stage. While these viewpoints give critical insights that can be correlated with the different stages of the disease in MS patients, the inclusion of intermediate timepoints might allow further granularity and uncovering of additional cellular states in MOLs.

The observed priming of OL differentiation and myelination gene programs in MOL5/6 particularly at the early stages of EAE suggest that there might be a window of opportunity for the promotion of MOL-associated remyelination in MS. Nevertheless, our results indicate only potential given the nature of epigenomic data and it is thus unclear whether it is possible to harness this MOL capability to promote remyelination. A deeper analysis of the epigenetic landscape at a single-cell level, by examining, for instance, activating and repressive histone modifications^54^or DNA methylation, might further elucidate mechanisms that could lead to the activation of these gene programes and promote remyelination in the context of MS.

## Methods

### Animals

Sox10:Cre-RCE:LoxP (EGFP) transgenic mice were used in this study. Sox10:Cre-RCE:LoxP (EGFP) mice are a strain of mice obtained originally by crossing mice with Cre recombinase under the control of the *Sox10* promoter (The Jackson Laboratories 025807; with a C57BL/6 genetic background) with reporter mice RCE:loxP-EGFP (with CD1 background, 32037-JAX) to label the complete oligodendrocyte (OL) lineage. Breedings were carried out with Cre allele females and non-CRE carrier males, with 1 male and up to 2 females. Breeding males carrying a hemizygous Cre allele, along with the reporter allele, with non-Cre females was avoided since it resulted in progeny expressing EGFP in all cells. Mice included in the experiment were heterozygote and between 9 and 13 weeks old, both male and female were included. Mouse housing conditions: dawn 6:00–7:00; daylight 07:00–18:00; dusk 18:00–19:00; night 19:00–06:00. A maximum of 5 adult mice per individually ventilated cage of type II (IVC seal safe GM500, tecniplast). All animals were free from mouse viral pathogens, ectoparasites and endoparasites and mouse bacteria pathogens. General housing parameters such as relative humidity, temperature, and ventilation follow the European Convention for the Protection of Vertebrate Animals used for experimental and other scientific purposes treaty ETS 123, Strasbourg 18.03.1996/01.01.1991. Briefly, consistent relative air humidity of 55% ± 10, 22 °C and the air quality is controlled with the use of stand-alone air handling units supplemented with HEPA filtrated air. Monitoring of husbandry parameters is done using ScanClime® (Scanbur) units. Cages contained hardwood bedding (TAPVEI, Estonia), nesting material, shredded paper, gnawing sticks and card box shelter (Scanbur). The mice received a regular chow diet (either R70 diet or R34, Lantmännen Lantbruk, Sweden). Water was provided by using a water bottle, which was changed weekly. Cages were changed once every week. All cage changes were done in a laminar air-flow cabinet. Facility personnel wore dedicated scrubs, socks and shoes. Respiratory masks were used when working outside of the laminar air-flow cabinet.

All experimental procedures on animals were performed following the European Directive 2010/63/EU, local Swedish directive L150/SJVFS/2019:9, Saknr L150, and Karolinska Institutet complementary guidelines for procurement and use of laboratory animals, Dnr. 1937/03-640. The procedures described were approved by the local committee for ethical experiments on laboratory animals in Sweden (Stockholms Norra Djurförsöksetiska nämnd), license numbers: 1995-2019 and 7029-2020.

### EAE

For the EAE induction, animals were injected subcutaneously with an emulsion of MOG_35–55_ in complete Freund’s adjuvant (CFA) (Hooke Laboratories, EK-2110, containing 1 mg MOG35-55/mL emulsion and 2-5 mg killed mycobacterium tuberculosis H37Ra/mL emulsion, Hooke Laboratories adjusted concentrations by lot) followed by the intraperitoneal injection of pertussis toxin (Hooke Laboratories, included in the induction kits) 1x PBS (Gibco, 10010023) on day 0 and day 1 (200-225 ng per animal, adjusted by lot according to Hooke manufacturer’s instructions). Scores of EAE were graded according to the following criteria: 0, asymptomatic; 1, limp tail or titubation; 2, limp tail and weakness of hindlimbs; 3, limp tail and complete paralysis of hindlimbs; 4, limp tail, complete paralysis of two hindlimbs with forelimb involvement; 5, moribund or dead; 0.5 for intermediate symptoms.

CFA-control mice were injected subcutaneously with the control emulsion containing CFA but without MOG_35–55_ (Hooke Laboratories, CK-2110), followed by the administration of pertussis toxin in PBS on day 0 and day 1 (200-225 ng per animal, adjusted by lot according to Hooke manufacturer’s instructions). Spinal cords from EAE and CFA-Ctrl were collected at the early (days 8-9), peak (days 14-15), and late stage (days 38-40). Naïve-Ctrl spinal cords were collected from 3-month-old mice from the same strain. Scores for EAE and CFA-Ctrl were plotted using GraphPad Prism version 9.0.0.

### Tissue dissociation, FACS sorting, and single cell multiome RNA-seq + ATAC-seq

At the early, peak, and late stage, mice were perfused with PBS, and spinal cords were collected. Spinal cord tissues were then dissociated into a single cell suspension according to the manufacturer’s protocol of Adult Brain Dissociation Kit, mouse and rat (Miltenyi Biotec, 130-107-677, we did not perform the red blood cells removal step since the majority of the red blood cells had been removed with PBS perfusion). For the debris removal step, the majority of the samples were processed with 38% percoll except 2 Naïve-Ctrl samples, 2 EAE peak stage samples, and 2 EAE late stage samples were processed debris removal solution (Miltenyi Biotec, 130-107-677) according to the manufacturer’s protocol to enrich more OPCs. Spinal cord single EGFP+ cells were enriched with the BD FACS Aria III Cell Sorter (BD Biosciences).

The cells were then lysed and washed according to the demonstrated protocol, Nuclei Isolation for Single Cell Multiome ATAC + Gene Expression Sequencing (10x Genomics, CG000365), with modifications: the cells were centrifuged for 10 min at 300 g and 4℃, resuspended in lysis buffer (containing 10 mM Tris-HCl pH 7.4, 10 mM NaCl, 3 mM MgCl2, 0.01% Tween-20, 0.01% IGEPAL (CA-630), 0.001% Digitonin, 1% BSA, 1 mM DTT, 1 U/ul RNase inhibitor) and incubated on ice for 3 min. After the incubation, wash buffer (containing 10 mM Tris-HCl pH 7.4, 10 mM NaCl, 3 mM MgCl2, 0.1% Tween-20, 1% BSA, 1 mM DTT, 1 U/ul RNase inhibitor) was added on top without mixing. The nuclei were centrifuged for 5 min at 500 g and 4℃. Nuclei were washed once in wash buffer followed by another wash with diluted 1x Nuclei buffer (10x Genomics, PN-1000283) containing 1% BSA, 1 mM DTT, and 1 U/ul RNase inhibitor. After that, the Chromium Next GEM Single Cell Multiome ATAC + Gene Expression chemistry (10x Genomics, PN-1000283) was applied to create single cells ATAC and RNA libraries. 26 EAE mice (8 mice from early stage, 8 mice from peak stage and 10 mice from late stage), 12 CFA-Ctrl mice (4 mice from early stage, 4 mice from peak stage and 4 mice from late stage), and 4 naïve-Ctrl were used for independent replicates. Libraries were sequenced on an Illumina Novaseq 6000 with a 50-8-24-49 read setup for ATAC (minimum 25,000 read pairs per cell) and a 28-10-10-90 read setup for RNA (minimum 20,000 read pairs per cell).

Both male and female mice were used in our study. The majority of the samples contain cells from one male mouse and one female mouse, except 2 samples from EAE early stage (one only contains male mice, another only contains female mice), 2 samples from the EAE peak stage (one only contains male mice, another only contains female mice), and 2 samples from the EAE late stage (one only contains male mice, another only contains female mice) for establishing sex prediction model and validation.

### RNAscope and confocal microscopy

RNAscope ISH was performed on 14μm spinal cord sections from the lumbar spinal cord of controls and EAE mice (n = 3 for each condition) with probes for mouse *Sox10* (ACD, 435931) and *H2-Ab1* (ACD, 414731-C2). RNAscope ISH protocol for sections was performed following the manufacturer’s protocol with minor modifications (RNAscope® Multiplex Fluorescent Detection Reagents v2, 323110): tissue sections were incubated in 1x target retrieval (RNAscope® Target Retrieval Reagents, 322000) for 5 min at 98℃ followed by 2 times of 2 min washes with DNase/RNase-free water. Incubated with protease IV (RNAscope® Protease III & Protease IV Reagents, 322340) at room temperature for 20 min followed by 2 times of 2 min washes with DNase/RNase-free water. Hybridized with probes (C1:C2/C3 in 1:50 dilution) for 2 h at 40 °C and washed twice with wash buffer (RNAscope® Wash Buffer Reagents, 310091). Amplification steps were performed by incubating with v2Amp1 (30 min), v2Amp2 (30 min) and v2Amp3 (15 min) (RNAscope® Multiplex Fluorescent Detection Reagents v2, 323110) at 40 °C with 2 times of 2 min washes with wash buffer in between steps. Incubated the sections with v2-HRP-C1 for 15 min at 40°C, after 2 times of 2 min washes with wash buffer, add TSA-conjugated fluorophores (1:1500 dilution in TSA buffer (RNAscope® Multiplex TSA Buffer, 322810)) on the sections and incubated for 30 min at 40 °C. Wash the sections 2 times with wash buffer, and incubate with HRP blocker for 30 min at 40 °C. Repeat the v2-HRP, TSA-conjugated fluorophores and HRP blocker steps for C2 and C3 channels subsequently if applicable. At the end, incubated with DAPI (Sigma, D9542, 1:5,000) for 5 min and washed with PBS with 0.05% Tween-20 (VWR, 9005-64-5) for 2 min.

Images of RNAscope were acquired using a Zeiss LSM800 Confocal, and processed in Fiji/ImageJ. Six 20X randomly selected fields per mouse (three from lesion and three from non-lesion) were chosen for quantification. Two-way ANOVA with Tukey’s multiple comparisons test was performed using GraphPad Prism version 9.0.0 for comparing the percentages of Sox+ H2-Ab1+ cells between different time points and lesion vs non-lesion.

### Raw data processing

A total of 7 batches were collected from the sequencing facility. Fastq files from 21 samples were processed throughout the 10X genomics standard pipeline. Gene expression and chromatin accessibility libraries were inputed into cellranger-arc2^55^ ‘count’ v2.0.2 with default settings to align the biological readouts on the associated mm10 reference genome v2020-A-2.0.0. Sample aggregation of both transcriptomic and genomics metrics was done using the ‘aggr’ of the same cellranger executable file, without normalization.

The aggregated count matrix and fragments file were loaded into R v4.2.3 (https://www.R-project.org). The former was lodged into a Seurat v4.3.0^56^ assay while the latter were accommodated into a Signac v1.9.0^57^chromatin assay associated to an Ensembl^58^base annotation v79 for mice where UCSC nomenclature was applied to provide gene names readability.

If not specifically mentioned, the following depictions of the analysis pipeline belong to either Seurat or Signac packages.

After assessing that each cell could be uniquely identified, quality metrics for both gene expression and peak accessibility were calculated. Among others, the mitochondrial ratio, the cell cycle score, the nucleosome signal, and the Transcription Start Site (TSS) enrichment were generated to support the following quality control cutoffs.

Depending on the quality of each sample, for each cell, a maximum of 30,000/150,000 and a minimum of 1,000 ATAC counts, a maximum of 10,000/45,000 and a minimum of 600 RNA counts, a minimum of 600 detected genes, a maximum of 1/1.5 nucleosomal signal, a TSS minimum enrichment of 2 and a maximum percentage of mitochondrial information of 15/25 were prerequisites to consider a given cell for the downstream analysis. These thresholds shrank by around 12% the overall number of cells, from 104,479 down to 91,757. While these thresholds are on purpose not too strained, they did not lead to any clustering perturbation downstream.

### Peak calling

The aggregated fragments file was used in MACS2 v2.2.6^59^to proceed to a peak call. Peaks falling in non-standard chromosomes or overlapping genomic blacklist regions from the mm10 genome were discarded. The Fraction of Reads in Peaks (FRiP) was then calculated for each cell, and the mean value for each sample ranged from 0.74 to 0.81.

### Gene activity

The computation of fragment counts per cell in the gene body and promoter region (Gene activity) was done with the GeneActivity function from Signac fetching all biotypes and extending 500bp upstream the TSS to catch the promoter region. All other parameters were set to default. This matrix was used later on to infer chromatin accessibility for each gene.

### Doublet detection

The detection of putative events where more than one cell in the same droplet occurs was done using DoubletFinder v2.0.3^60^. The gene counts matrix for individual sample was normalized by library size, multiplied by 10,000 and natural-log transformed. The expression of the top 1,000 most variable genes was centered by subtracting their average expression, scaled by dividing their standard deviation, and used for a Principal Component Analysis (PCA). The Euclidean distances between each cell were calculated using the first 15 Principal Components (PCs) to build a Shared k-Nearest Neighbors (SNN) graph. A Louvain algorithm was used to optimize the modularity of the SNN graph to determine clusters at a resolution of 0.5. Then a Uniform Manifold Approximation and Projection (UMAP) dimensional reduction was applied to the first 15 PCs to project the cells on a 2D space.

The pN-pK parameter sweep was performed with the paramSweep_v3 function on the first 15 PCs to a maximum of 10,000 cells and then pass on the summarize Sweep function to summarize and compute the bimodality coefficient across pN and pK parameter space. The mean-variance normalized of the bimodality coefficient was computed using the find.pK function to highlight the optimum pK value of each sample. The best pK value of each sample was fluctuating between 0.005 and 0.7. While the number of generated artificial doublets (pN) was set to 0.25 as a default parameter for all samples, the prediction of the proportion of the total number of doublets (nExp) per sample was estimated by creating a model of the proportion of homotypic doublets via the annotated clusters mentioned above. The homotypic proportion complement was then multiplied by the number of cells in the sample times the best pK of the same sample. The first 15 PCs, the pN, best pK, and nExp were set as parameters in the doubletFinder_v3 function to calculate the proportion of Artificial Nearest Neighbors (pANN) for each real cell. This pANN score is then compared to the number of expected doublets to generate the final doublet predictions.

Almost all samples drawn less than 1% of doublets but 4 samples drawn 1.47%, 3.32%, 5.93% and 11.33% of doublets. Using the multiplet rate provided by 10X genomics (https://kb.10xgenomics.com/hc/en-us/articles/360001378811-What-is-the-maximum-number-of-cells-that-can-be-profiled-) we were able to measure a 0.8%/7.2% theoretical number of doublets. Predicted doublets were removed from the downstream analysis.

### Sex classification model

A total of 30,749 cells were sequenced with a priori sex annotation (18,245 cells were female and 12,504 cells were male) from EAE time points (7302 cells from early, 8214 cells from peak and 15,233 cells from late stage). This knowledge was used to create a machine-learning sex classifier to investigate sex transcriptomic or epigenetic differences. An equal number of cells for each gender (12,504 cells) were extracted from the main object to a sex object to build the model. Mouse genes and positions from chrX and chrY were extracted from Ensembl Mart v2.54.1^61^. For each chromosome and each cell, the sum of the gene count expression of those existing in the sex object was divided by the total number of counts by cell, generating a percentage of reads falling into transcribable regions on either chrX or chrY. In addition, a gene score was calculated from *Xist* using the AddModuleScore from Seurat, with 200 control features from the same bin per analyzed feature. The sex object was then again subdivided into a training dataset and a testing dataset, representing 80% and 20% of the dataset, respectively. In association with the sex annotation, these three scores were sufficient to train a random forest classifier from the Caret v6.0 package in R, able to differentiate male and female cells with high accuracy.

The random forest function was run using the train function in Caret. The trControl parameter, for controlling the computational nuances of the main function, was calculated using the trainControl function with “cv” method, 5 number of folds, derivating from the createFolds function on the sex annotation to 5 folds, asking for a “grid” search and to class the probabilities turning on the “classProbs” parameter, saving the final predictions in two classes. The tuneGrib parameter was set to consider the number of variables to randomly sample (mtry) as 1 and 2. The model was trained sequentially on different node sizes (20, 30 and 50) and number of trees (5,000, 10,000 and 20,000). The remaining parameters were set as default and the ROC value calculation was turned on to assess the best node size, number of trees and mtry combo. Not by far, the association of 20,000 trees and 20 nodes and 1 mtry outperformed the other combinations.

The validation on the testing dataset yielded an accuracy of 95.3%, a sensitivity of 91.0% and a specificity of 99.7%. An insignificant number of male cells were annotated as female cells (accuracy of 99.48%), while a few female cells were assigned to male cells (accuracy of 86.13%). We then used the newly created model to process each cell of the main object. Correction of miss assigned cells (8.4%) from a priori sex annotated samples was performed.

### Normalization

Each feature of each cell of the gene count matrix was divided by the total counts of that cell and multiplied by a scale factor of 10,000. The obtained scaled matrix was then natural-log transformed. The peak count matrix was processed through a Term Frequency Inverse Document Frequency (TF-IDF) normalization, which computed a log(TF x IDF) with a scale factor of 10,000. Each feature of each cell of the gene activity matrix was divided by the total counts of that cell and multiplied by a scale factor of 10,000. The obtained scaled matrix was then natural-log transformed.

### Significant features selection

In order to proceed to the reduction of the dataset dimensions, the 2000 most variable genes were selected via variance stabilizing transformation, and 95% of the most common peaks were also selected. The expression of each gene was transformed by subtracting its average expression and scaled by dividing its standard deviation.

### Dimension reduction

The reduction of the dimensionality of the expression matrix on the 2,000 most variable genes was done by running a Principal Component Analysis (PCA). The first 30 Principal Components (PCs), representing most of the diversity of the dataset, were selected by the elbow plot method. The partial Singular Value Decomposition (SVD) was used to reduce the dimensions of the most common peaks and 25 of the first Latent Semantic Indexing (LSI) layers were selected via elbow method. However, the first LSI was removed from the selected dimensions due to its high correlation with chromatin capture depth, which would bias the investigation of the chromatin opening binarity.

### Construction of the nearest-neighbor graph

For both, principal components and latent semantic indexing, the construction of the graphs were made on “annoy” method with Euclidean distances and 50 trees. A “k” for the k-nearest neighbor algorithm was set to 20. For the construction of the graph based on principal components, an acceptable Jaccard index was set to 0. For the multi-modal nearest-neighbor graph, the same number of PCs and LSIs were used. A “k” for the k-nearest neighbor algorithm was set to 20, 200 approximate neighbors were computed and the cutoff to discard the edge in the SNN graph was set to 0.

### Cluster determination

The unsupervised clustering of the nearest-neighbor graphs was done by the Louvain algorithm. The resolution of each clustering was assessed using the “clustree” R package iterating over different resolutions to pinpoint the most stable clustering resolution. Resolutions of 2, 1.5 and 1.7 were selected for gene expression, peak accessibility and joined modality, respectively.

### Cell projection

In allow visualization of the cells despite the high dimensionality of the data, the nearest-neighbor graphs were reduced to only two dimensions using UMAP with the appropriate number of PCs and LSIs (See Dimension Reduction section), and other values set as default.

### Batch effect correction

As sample libraries were not prepared on the same day and sequenced in the same sequencing run, technical artifacts might arise and generate variability not related to biology in the dataset. However, accounting for such disequilibrium, without removing the subtle differences between timepoints was challenging. Batch effect correction methods such as Harmony^62^or Scanorama^63^with soft parameters totally masked the differences between timepoints. Moreover, integration of the sample with “cca” and “rpca” was also over-correcting the time point-specific cell populations (data not shown). Although the popular batch effect methods were not suitable for our peculiar dataset, we rely on the strength that every sequencing run possesses at least one CFA-Ctrl sample (except on late time point and Naïve-Ctrl samples). Thus, we investigate if the cells in each CFA-Ctrl cluster were well-mixed across the different replicates. All 14 major CFA-Ctrl clusters contain cells from all replicates. To go further at the cell mixing level, a Local Inverse Simpson’s Index (LISI) from Harmony, accounting for categorical variable diversity, was calculated for an equivalent number of cells in each CFA-Ctrl replicate, using the first two components of the joint projected UMAP. This score, initially ranging from 1 to the number of replicas, has been normalized to a scale of 0 to 1. For CFA-Ctrl, 57% of the cells maintained an LISI of at least 0.5. The same strategy was applied to the Naïve-Ctrl samples where 15 of the 17 majors Naïve-Ctrl clusters contained mainly cells ascending from the two replicates, one homogeneous cluster was coming from a microglia cluster, and the other one was from a pericyte population. At the single-cell level, 80% of the cells maintained an LISI of at least 0.5. For all EAE time points, the same method was applied, yet the calculated LISI was lower than CFA-Ctrl or Naïve-Ctrl (EAE score equal to 0), probably due to the heterogeneity of the EAE score at the sample collection within each time point. Even though this method does not allow us to mitigate potential hidden batch effects, it grants us an overview of the satisfying cell distribution at a consistent EAE score.

### Label transfer

After removing cells with less than 200 detected genes, more than 8000 detected genes and less than 10% of reads matching the mitochondrial genome, from a literature dataset^5^, the transcriptome of the remaining cells was normalized and scaled using SCTransform from Seurat. A label transfer was then performed from the literature annotation to the present dataset, based on the transcriptional profile of selected anchors via FindTransferAnchors function using Canonical Correlation Analysis (CCA), the 10 first dimensions of the PCA, and recomputing the residual with the reference SCT model parameters. The label prediction was run using TransferData with the previously determined anchors set and the CCA reduction to weight the anchors. For each cell, the highest prediction score was selected to make the annotation.

### Cell annotation

Cell type annotation was assessed per cluster on the joined graph clustering resolution, both label transfer outputs, cell type marker gene expression and chromatin accessibility were taken into consideration. More refined cell types for the MOL population were later considered.

### Immune score

Exhaustive lists of immune-related genes published in Meijer &Agirre et al., Neuron 2022^12^, specific to “immune response” (GO:0002250) and “immune system process” (GO:0002376), were used to evaluate the immune capability of each cell. Cells from Naïve-Ctrl were used as the background signal. After removing unexpressed genes in our dataset, sub-categories with genes related to positive regulation of immune response, negative regulation of immune response, antigen processing and presentation, positive regulation of type I/II interferon-mediated signaling pathway, positive regulation cytokine production, complement activation, CD4^+^ T cell-related immune response and CD8^+^ T cell-related immune response were extracted from the immune gene list respectively, using the MGI immune database (https://www.informatics.jax.org/vocab/gene_ontology).

For each cell type, immune scores were calculated for each immune sub-category and as a whole, using the AddModuleScore function from Seurat with default parameters. All scores were scaled from 0 to 1, and the probability of each cell following the immune score distribution of the Naïve-Ctrl population, which followed a normal distribution, were processed using the following distribution function, where *μ* and *σ* are the mean and the standard deviation of the immune score distribution of the Naïve-Ctrl population, respectively:

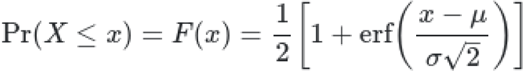

Under the null hypothesis that each cell follows the immune score distribution of the Naïve-Ctrl cells in a given population, each cell with a p-value<= 0.05, rejecting the null hypothesis, was classified as “Immune”. Respectively, the cells accepting the alternative hypothesis (pvalue> 0.05), sharing a relationship between the immune score of a given cell and the immune score distribution of the Naïve-Ctrl population, were classified as “Non Immune”. Due to the lack of cells annotated as Neurons in the Naïve-Ctrl samples, this population was not annotated.

The same approach was used on the gene activity matrix to compare transcriptional and epigenetic immune states.

### Naïve-Ctrl vs. CFA-Ctrl comparison

In order to acknowledge the impact of the CFA and pertussis toxin (PTX) in CFA-Ctrl compared to Naïve-Ctrl, we first did a Pearson correlation on the gene expression matrix in each cell type between replicates of each time point. Then, a differential gene expression analysis was performed using a Wilcoxon rank sum test between the two time points in each cell type. Genes with pvalue adjusted lower than 0.05 with a log fold change threshold over 0.5 (calculated with FindMarkers function from Seurat) and at least expressed in 10% of the cells in at least one condition were selected for plotting.

We compared the difference between different CFA-Ctrl replicates and found high similarity of expression between OLs from different time points (Pearson correlation score between different CFA-Ctrl replicates: 0.95-0.99). Therefore, we pooled CFA-Ctrl from different time points together for the subsequent comparison with EAE.

In microglia (MG), however, we found the expression of some genes had significant differences between replicates, such as Il1a, Atf3, and Ier5 (Pearson correlation score between different CFA-Ctrl replicates: 0.69-0.90, **Extended Data Fig. 10a**). Since the cells we applied for sequencing were sorted based on GFP signal, these MG are likely to be activated MG that engulf OL, therefore more sensitive to the ex vivo alteration. The finding that MG are more sensitive than other CNS cell types has also been confirmed before in another study^64^. We did not compare the difference between CFA-Ctrl replicates in neuron, ependymal cells, and pericytes due to the low cell number.

Since CFA-Ctrl was also admitted to the subcutaneous injection of emulsion containing CFA and intraperitoneal injection of PTX, which creates an immune response driven increase of blood-brain barrier permeability, we compared the gene expression between CFA-Ctrl and Naïve-Ctrl. We found that genes related to metabolic processes, such as mt-Atp6, mt-Cytb, and my-Co3, had higher expression in both MOL2 and MOL5/6 from Naïve-Ctrl than CFA-Ctrl. Ribosomal genes, such as Rps10, Rpl18, and Eif1 were increased in MOL2 and MOL5/6 from CFA-Ctrl compared to Naïve-Ctrl (**Extended Data Fig. 10b**). However, when comparing MG between Naïve-Ctrl and CFA-Ctrl, we observed a notable enrichment of MG activation and immune system process related genes in CFA-Ctrl compared to Naïve-Ctrl, such as H2-K1, Mt1, and Cd68 (**Extended Data Fig. 10c,d**). Additionally, genes associated with translation, such as Rps27a, Rpl26, and Eif1, were also found to have increased expression in MG from CFA-Ctrl (**Extended Data Fig. 10c**). These findings suggest that the administration of CFA and PTX resulted in the activation and increased proliferation of MG, although it did not trigger myelin-specific immune response.

### OL lineage investigation

All cells belonging to the OL lineage were subsetted for fine-tuning annotation. One EAE early time point sample collected on day 8 post-immunization with a score of 0 (without any symptom but with weight loss during the disease course) was removed from the analysis due to no EAE symptom and similar gene expression as CFA-Ctrl. As no major gene expression differences were found between Naïve-Ctrl and CFA-Ctrl in OL lineage population, Naïve-Ctrl samples were removed from the downstream analysis. The processing of this OL lineage subset of 68499 cells, is similar to the one with all cell types. Nevertheless, few differences have to be mentioned, only 1000 genes were selected for the PCA and the 35 first PCs and the first 20 LSIs were selected for graphs construction. Last, cluster resolutions 2.2, 0.9 and 3.3 were selected for gene expression, peak accessibility and joined modality respectively. An in-depth label transfer was performed on the subset with the same methods as previously described, using only the OL lineage cells from the literature dataset^5^**(Extended Data Fig. 3c,d)**. Unsupervised aggregation of clusters was assessed on joined graph clustering resolution, after calculation of Pearson correlation on log2 gene expression and computation of silhouette information for different number k of clusters. The determination of k was set to the first value generating only positive silhouette width values after proper splitting of the main OL lineage cell types (OPC, COP, MOL1, MOL2, MOL56), k=29. Silhouette values for k-medoids clustering were calculated with the “pam” function from the R package “cluster” v.2.1.4. OL lineage cell type sub clustering (α, β, γ, δ, ε, ζ, η, θ, ι, κ, λ, μ, ν) was assessed per aggregated cluster and ordered along their average immune score for each main OL lineage cell type **(Extended Data Fig. 4a)**.

### Multimodality features selection

Transcriptomic and epigenomic variations in specific cell types throughout the disease time course were performed in a pseudo bulk manner. This technique allows us to overcome the sparsity of the datasets and highlight dynamic features within 3 modalities, gene expression, gene/promoter accessibility, and peak accessibility. These tri omics modalities meet the prerequisites for analyses based on the negative binomial distribution (lack of intra sample variability, most of the features remain stable throughout the tested conditions, raw integer counts to specific genomic locations, sufficient number of replicates). Each modality was loaded into a SingleCellExperiment v1.20.1^65^object where features with less than 10 counts were removed. Feature counts were then aggregated across cell types and broken down by sample. Each cell type was processed independently, samples formed by less than 5 cells and time points formed by less than 30 cells were not considered for the downstream investigation. The resulting matrix and associated metadata were loaded into a R object using DESeq2 v1.38.3^66^, with a design formula including sample names in addition to the debris removal method as a covariate. After size factors and dispersion estimations, results tables were extracted per contrast. We decided to investigate 4 contrasts (Early/Ctrl, Peak/Early, Late/Early, and Peak/Late), depicting a broad overview of the disease time course. Features presenting an absolute log2 fold change over 1 with a pvalue adjusted lower than 0.01 and a baseMean (the average of the normalized count values, dividing by size factors, taken over all samples) superior to 1, were selected to be displayed on the heatmaps and gene ontology analyses for the modalities represented by gene and promoter, and included into the transcription factor analysis for the modality represented by peaks. The differential gene expression between male and female within each time point was performed similarly, merging gene counts by both sample and sex. The debris removal method was included in the differential expression formula as a covariate for Peak and Late time points.

### Multimodality pseudotime

The expression dynamic was assessed by velocyto v.0.17.17^26^to retrieve the number of spliced and unspliced reads in each sample. In addition to the cellranger output directory, the gtf annotation file used by cellranger was given, along with a gtf annotation file for mm10 repetitive elements from RepeatMasker (https://www.repeatmasker.org). Resulting loom files were loaded into scVelo^67^in python, and directly imported via MultiVelo v.0.1.2. After merging the samples, OL lineage cells were selected to undergo a log-transformed normalization on the 1,000 most variable features with at least 10 read counts.

The chromatin accessibility matrices from cellranger outputs were loaded and peaks were aggregated around each gene and those in correlation with promoter peak or gene expression as well, for a maximum distance of 10,000 bp, using the “feature linkage” prediction from cellranger outputs. After merging the samples, OL lineage cells were selected to undergo a TF-IDF normalization.

The 68,499 cells as well as 547 genes with both expression and chromatin accessibility signals were picked to compute moments for velocity estimation for each cell across its 50 nearest neighbors calculated from Euclidean distances in the first 30 PCs space of the expression matrix. Weighted nearest neighbor properties calculated previously on the OL lineage cells were used to smooth the epigenetic modality and be incorporated into RNA velocity to recover chromatin dynamics and carry out enhanced lineage predictions. This last step was done on MOL56 and MOL2 separately. For each cell type, a new UMAP was generated, velocity, latent time as well as terminal states were processed, and CFA-Ctrl cells were set as root cells.

MultiVelo classifies, if possible, each gene into two modules of biological dynamics. A first model (M1) where the chromatin starts closing before the ending of the transcription and a second one (M2) where the chromatin starts closing after the end of the transcription. Moreover, the coupled kinetics of the transcriptomic and epigenomic profiles can be used as leverage to predict a current cell state for a given gene. The priming state is considered when the chromatin is opening but no unspliced transcript has yet been detected. The couple-on state is selected when the chromatin is opened and the number of unspliced transcripts is increasing. The decoupling state is picked when there is a decorrelation between chromatin and unspliced transcripts dynamics. For the first model the chromatin closes before the end of the transcription, while for the second model the number of unspliced transcripts starts decreasing but the chromatin is still open. The couple-off state is set when the chromatin is closed while the number of unspliced transcripts is collapsing.

### Potential enhancers & Domains Of Regulatory Chromatin (DORCs)

The inter-modality investigation helped us associate detected peaks to the expression of the most probable nearby genes using LinkPeaks function from Signac, inspired by the method described in the SHARE-seq paper^24^. Each peak to gene connection, occurring in at least 10 cells and within a 50,000bp distance from the gene TSS, with a positive Pearson correlation coefficient and a pvalue lower than 0.05 was considered as a meaningful interaction. For each cell, the number of fragments falling into the 42,776 peaks involved in the same number of meaningful interactions was divided by the total number of fragments falling in peaks for the same cell. From the normalized matrix of meaningful peaks per cell, the normalized number of fragments in peaks were aggregated by gene using the peak to gene connections, for a total of 13,782 genes joined to at least one peak. The regulatory chromatin score for each cell of the generated genes was then multiplied by a scaling factor of 10,000. Notwithstanding the value of such enhancers score, we were interested in deciphering more consequent domains of regulatory chromatin. Therefore, we selected a pool of genes connected to at least 5 peaks to redo the peaks to genes connectivity calculation on a broader distance of 500,000bp from the gene TSS. Keeping positive Pearson correlation coefficient and a pvalue lower than 0.05, a total of 54,789 peaks were connected to a total of 2,818 genes defined as DORCs. Similarly, to the regulatory chromatin of genes, the DORCs were normalized, aggregated per gene according to the new connection material and scaled to generate a DORCs score.

As doing the sum of the peaks counts conserves the negative binomial distribution of the epigenetic modality, both regulatory chromatin and domains of regulatory chromatin regulated gene were reversed into unnormalized matrices (scaling factor and number of fragments in peaks for a given cell) to be loaded into a SingleCellExperiment object to ascertain, with confidence, gene variations between timepoints in a pseudo bulk manner as described above, with the same parameters and thresholds.

### Gene Regulatory Network (GRN) and predicted Transcription Factors (TF)

GRN was inferred with Pando 1.0.0^29^. Pando models the relationship between TFs and their binding sites in selected regulatory regions with the expression of target genes combining the multiome RNA and ATAC information.

The whole dataset was subset in MOL2 and MOL5/6, for each MOL we subset randomly a maximum of 2000 cells per time point. We selected peak regions specific to each time point in MOL2 and MOL5/6 (see in multimodality features selection) as candidate regulatory regions to scan for TF binding motifs, with JASPAR (2022 release).

The GRN was inferred by fitting Generalized Linear Models (GLM) implemented in Pando for the expression of each gene. The regression model with peak-TF pairs as independent variables and target genes as response variables was built for MOL2 and MOL5/6 independently. Peaks were assigned to nearby genes with the peak_to_gene_method=”Signac”. Which considers the closest distance, upstream or downstream, to the gene. The model estimates of each predicted TF with a target gene can be interpreted as a measure of interaction between both. We calculated a TF activity by multiplying the mean estimates (coefficient) in the model by the average TF expression at each time point. TF activities were ranked per highest positive activity in each time point. Gene modules were selected with a R2 threshold of adjusted p-value < 0.05 and default parameters with Pando find_modules function.

### Heatmaps

Samples with less than 5 cells and time points with less than 30 cells were removed from the creation of the heatmaps. For gene expression and gene/promoter accessibility, the log2 average raw gene expression and gene/promoter accessibility at each time point and cell type were calculated with an extra pseudocount to generate heatmaps scaled across time points from 0 (low) to 1 (high). For gene regulatory chromatin, the average value of the gene regulatory chromatin score at each time point and cell type was scaled across time points from 0 (low) to 1 (high). Missing values in any tested modalities were set to 0. The black column on the right side of each heatmap represents the gene average raw count. For each modality, the average of the previously calculated scaled values per gene category and time point was used to produce the summarized dynamic plots.

### Gene ontology

Highlights of biological pathways involving the most dynamic features along the disease time course were assessed using biomaRt v2.54.1 R package in addition to the Ensembl v2.22.0 database. For each cell type, the top 50 features with the highest amplitude across time points were selected to undergo a pathway enrichment via the “enrichPathway” function from ReactomePA v1.42.0 R package, with a pvalueCutoff set at 0.05, a qvalueCutoff set to 0.05, and false discovery rate (“fdr”) as a method of adjustment. From the same R package, dot plots were displayed by “GeneRatio” for 10 categories. Network graphs were also built from the “emapplot” function taking as input the result of the pairwise_termsim function, with Jaccard similarity coefficient as the calculation method for 50 categories.

For some specific categories, yielding no significant pathway enrichment (such as differentially expressed genes: Type IV in MOL2 and Type I in MOL5/6, differential chromatin accessibility of promoter + gene body: Type II in MOL2), the p-value threshold was removed. The generated gene ontology was then given as an indicator of potential enriched pathways.

### Genome tracks

The coverage of the DNA fragments within a given genomic region was done using CoveragePlot from Signac. Tracks were normalized by group using a scaling factor as the number of cells within the group multiplied by the sequencing depth average of the group.

### Single-cell genomic heatmaps

All fragments from 50 randomly selected cells per cluster and falling into a specific genomic region were carried into a 250bp windowed matrix that is further down binarized and plotted.

### Percentage of cells on stacked bar plots

The number of cells per sub-cell type in each condition was aggregated, normalized across conditions to get a proportion of each sub-cell type per condition and divided by the number of conditions in order to keep the sum of the proportion equal to 100%.

### Percentage of cells on side-by-side bar plots

The number of predicted male and female cells in each sex-specific sample was aggregated, and normalized across samples to get a proportion of each sex per sample.

### Circos plot

Number of cells in the top levels of the circos plots matching each bottom level of the circos plot. For an unbiased visualization of proportion, some circos plots are randomly downsampled by either top levels, down levels or both (details were mentioned in figure legends).

### Bigwig files

The mouse genome was segmented into 100bp windows and fragments were assigned to their corresponding window, generating a binarized genome per cell matrix. For each group of cells, the fragment counts matrix was aggregated, multiplied by a scaling factor of 10,000, and divided by the number of fragments in the group.

### Data availability

Raw fastq files, counts matrices and genomic tracks will be available on GEO. GEO deposition is ongoing and will be provided to the reviewers as soon as it is finalized.

### Code availability

Jupyter notebooks to process the raw datasets, as well as reproducing the figures are accessible at https://github.com/Castelo-Branco-lab/EAE_multiomics_2023/

## Acknowledgements

We thank Tony Jimenez-Beristain for writing the laboratory animal ethical permits 1995_2019 and 7029/2020 and for the assistance with animal experiments. We thank Elina Parvisto and Paria Samadi Tari at the Comparative Medicine Biomedicum facility (KI) for their advice and help with experimentation. We thank Maximilian Haeussler and Brittney Wick for preparing the UCSC Cell Browser entry for our dataset. We acknowledge support from the National Genomics Infrastructure in Stockholm funded by Science for Life Laboratory, the Knut and Alice Wallenberg Foundation and the Swedish Research Council. Part of the computation/data handling was enabled by resources provided by the National Academic Infrastructure for Supercomputing in Sweden (NAISS) and Swedish National Infrastructure for Computing (SNIC) at the Uppsala Multidisciplinary Center for Advanced Computational Science, partially funded by the Swedish Research Council through grant agreement no. 2022-06725 and no. 2018-05973. Part of the computing was also performed in the Linnarsson group Monod Linux cluster at MBB-KI, and we thank Peter Lönnerberg for maintenance and support. Chao Zheng was supported by the China Scholarship Council. Leslie Kirby was funded by an ECTRIMS postdoctoral fellowship. Work in G.C.-B.’s research group was supported by the Swedish Research Council (grant 2019-01360), the European Union (Horizon 2020 Research and Innovation Programme/ European Research Council Consolidator Grant EPIScOPE, grant agreement number 681893), the Swedish Brain Foundation (FO2023-0032), Knut and Alice Wallenberg Foundation (grants 2019-0107 and 2019-0089), the Göran Gustafsson Foundation for Research in Natural Sciences and Medicine, the Swedish Society for Medical Research (SSMF, grant JUB2019), Ming Wai Lau Centre for Reparative Medicine, and Karolinska Institutet.

## Notes

### Competing Interest Statement

The authors have declared no competing interest.

