## Extended Data Figures 1-11 for "Distinct transcriptomic and epigenomic responses of mature oligodendrocytes during disease progression in a mouse model of multiple sclerosis"

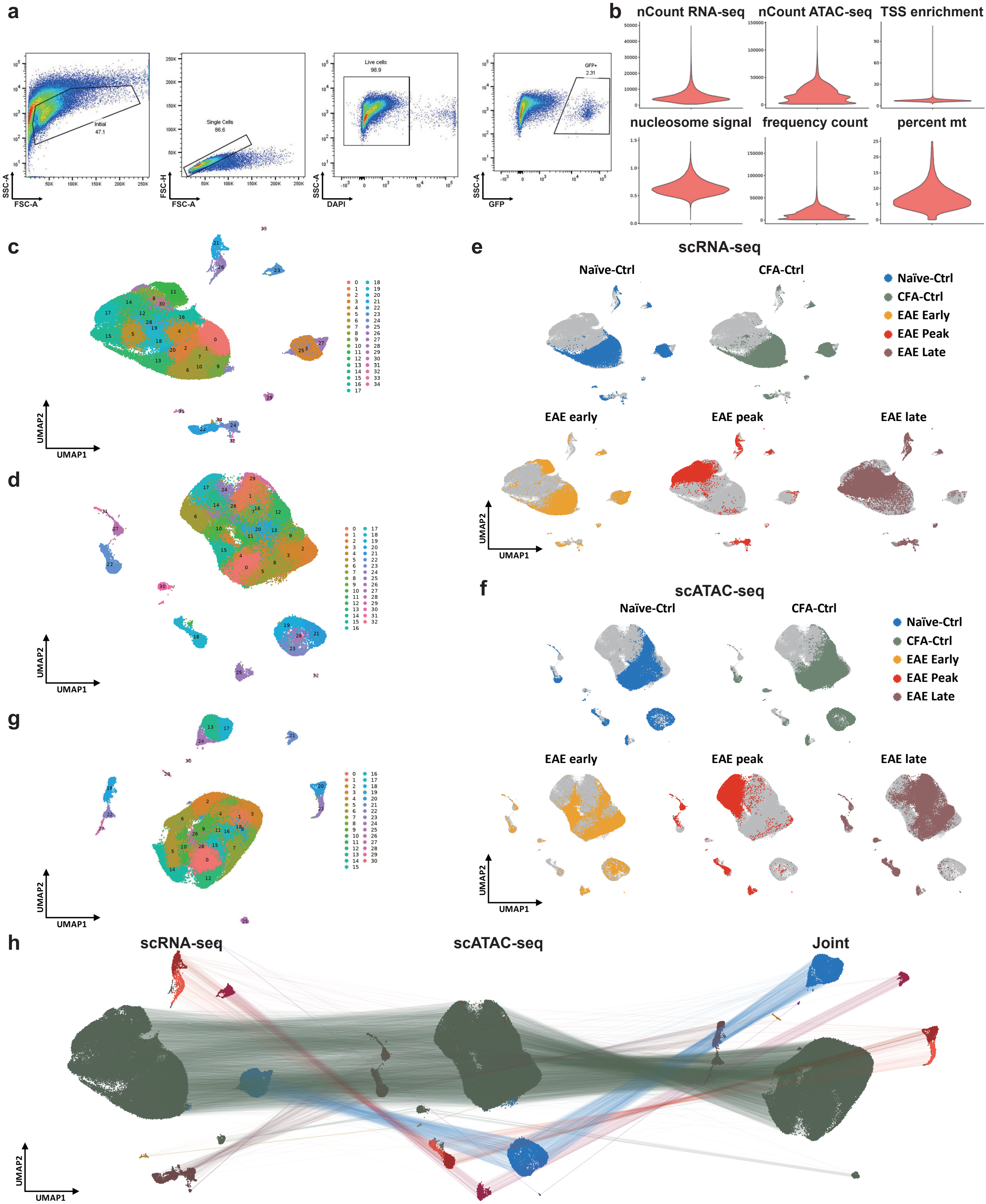


**Extended Data Fig. 1 Single cell multiome of OLGs from the EAE model.**
**a**, Representative fluorescence-activated cell sorting (FACS) gating strategy with a sample from EAE peak stage.
**b**, Quality control metrics after removing low quality cells of the multiome ATAC + gene expression sequencing data.
**c,d**, UMAP with Louvain clustering algorithm based on gene expression (c) and chromatin accessibility (d) of 10x Genomics chromium multiome ATAC + gene expression.
**e,f**, UMAP of the cells separated by condition based on gene expression data (e) and chromatin accessibility data (f).
**g**, UMAP with Louvain clustering algorithm based on joint projection of gene expression and chromatin accessibility modalities.
**h**, UMAP based on scRNA-seq data (left), scATAC-seq data (middle), and joint UMAP created by combining scRNA-seq and scATAC-seq nearest neighbors graphs (right), with lines connecting cells by single-cell barcodes across modalities.

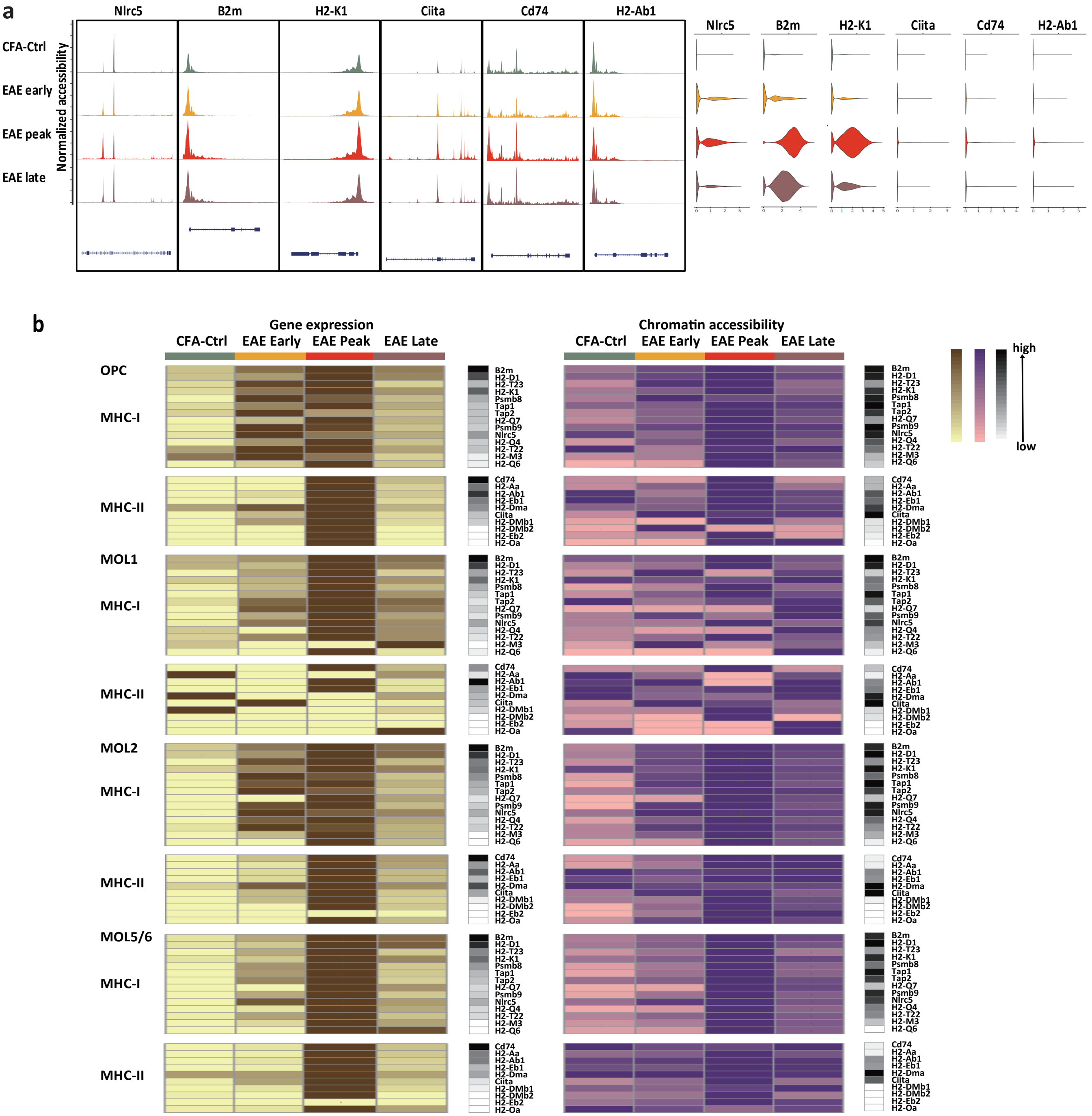


Extended Data Fig. 2 Expression and chromatin accessibility of MHC-I/II genes in different OLG cell types.
a, Normalized chromatin accessibility (left) and log2 expression (right) of representative MHC-I and II associated genes in each time point.
b, Heatmaps of the expression (left) and chromatin accessibility (right) of MHC-I and -II genes at different stages in OPC, MOL1, MOL2, and MOL5/6. The black column on the side represents the gene raw counts.

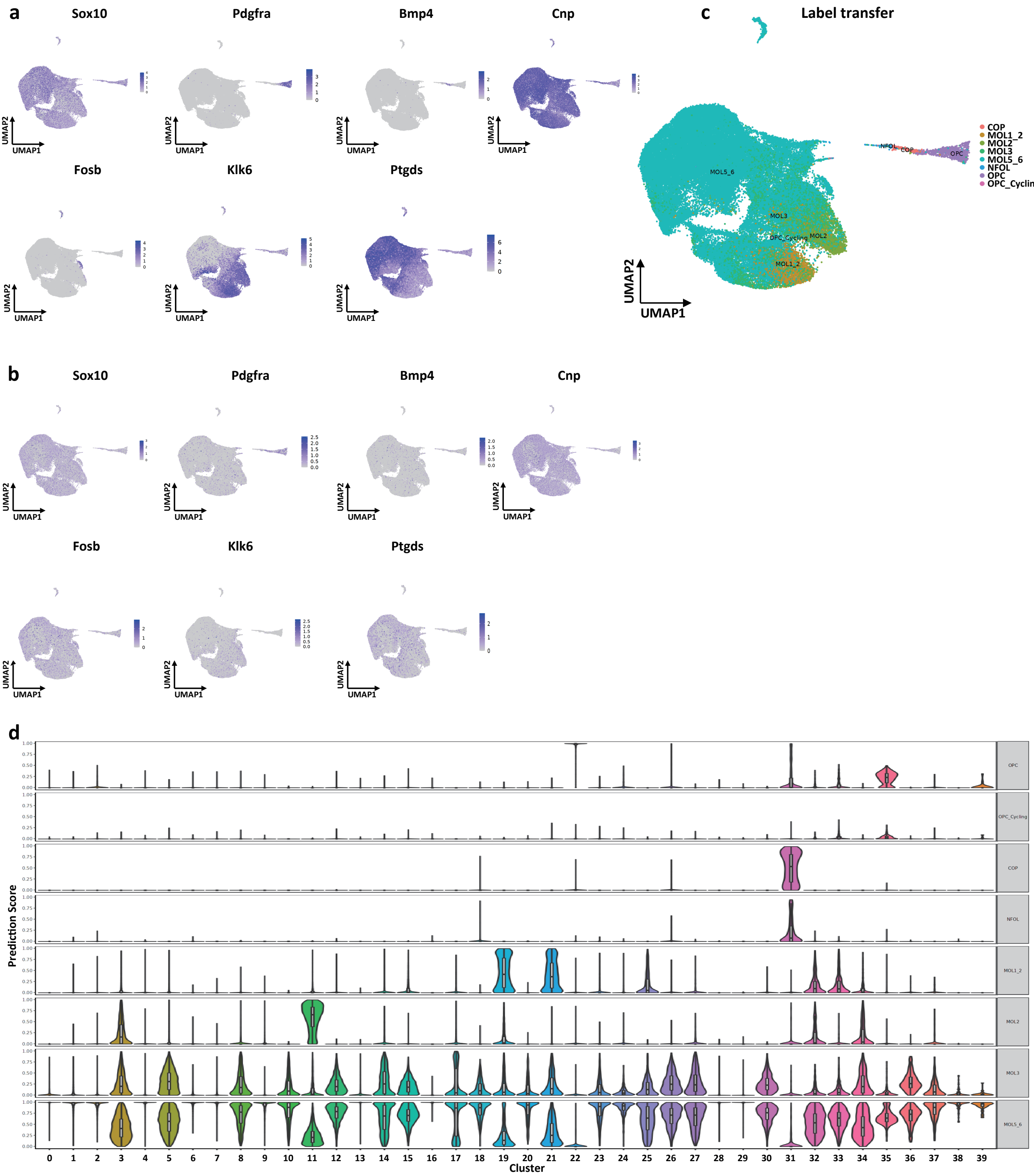


**Extended Data Fig.3 Cell type annotation based on gene expression and chromatin accessibility.**

**a,b, Feature plots showing gene expression (A) and chromatin accessibility activity score (B) of markers for OLG (Sox10), OPC (Pdgfra), COP (Bmp4), and MOL (Mbp, Fosb, Klk6, Ptgds) populations.**

**c, UMAP of label transfer predictions from matched scRNA-seq data.**

**d, Label transfer prediction scores from literature cell types to annotated clusters.**

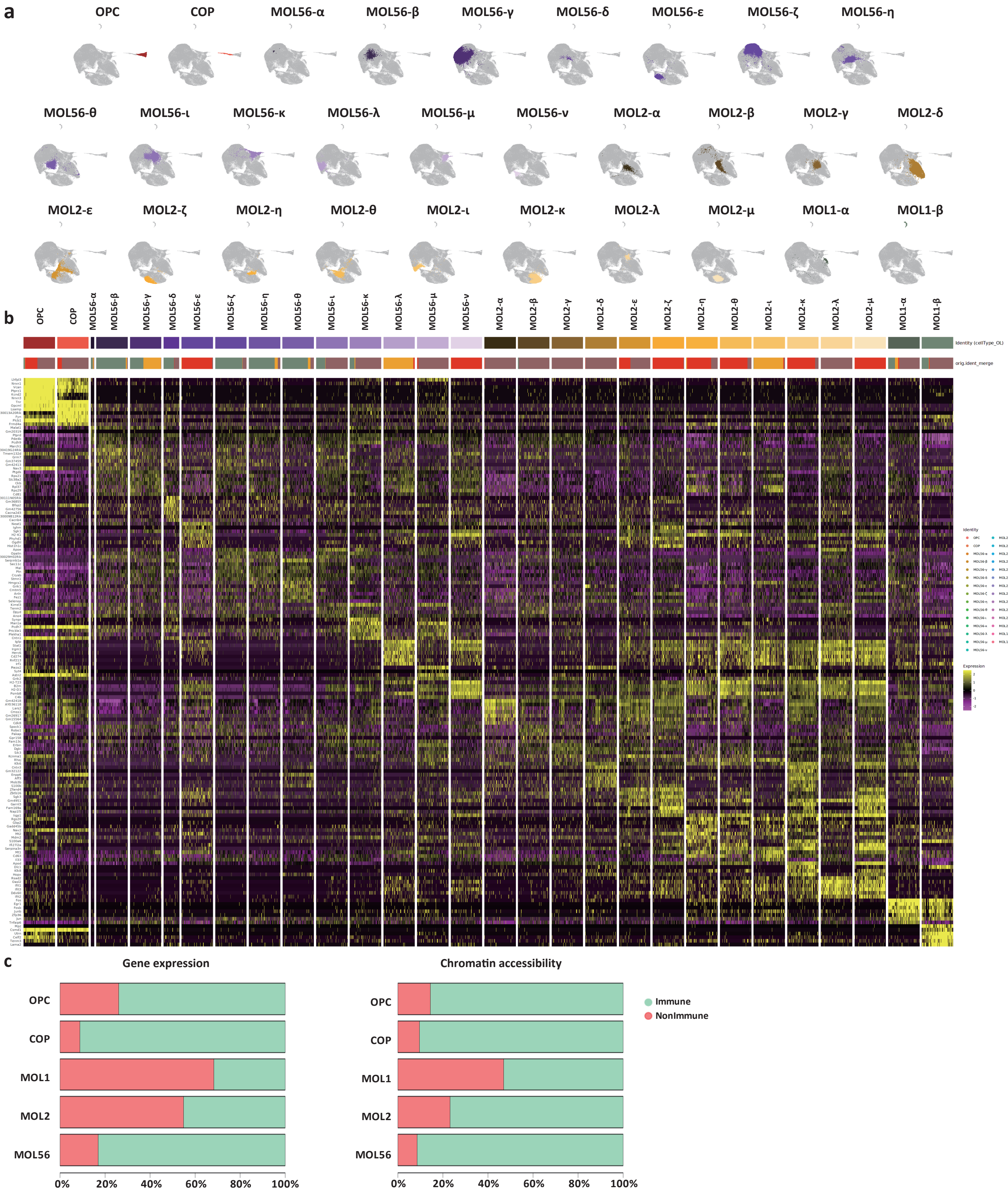


**Extended Data Fig. 4 MOL sub-cell type classification based on gene expression.**
a, Joint UMAP of OLG populations colored by sub-cell types.
b, Differentially expressed genes between OLG sub-cell types.
c, Percentage of cells with (red) or without (green) immune status identified by gene expression (left) and chromatin accessibility (right) in each OLG sub-cell type.

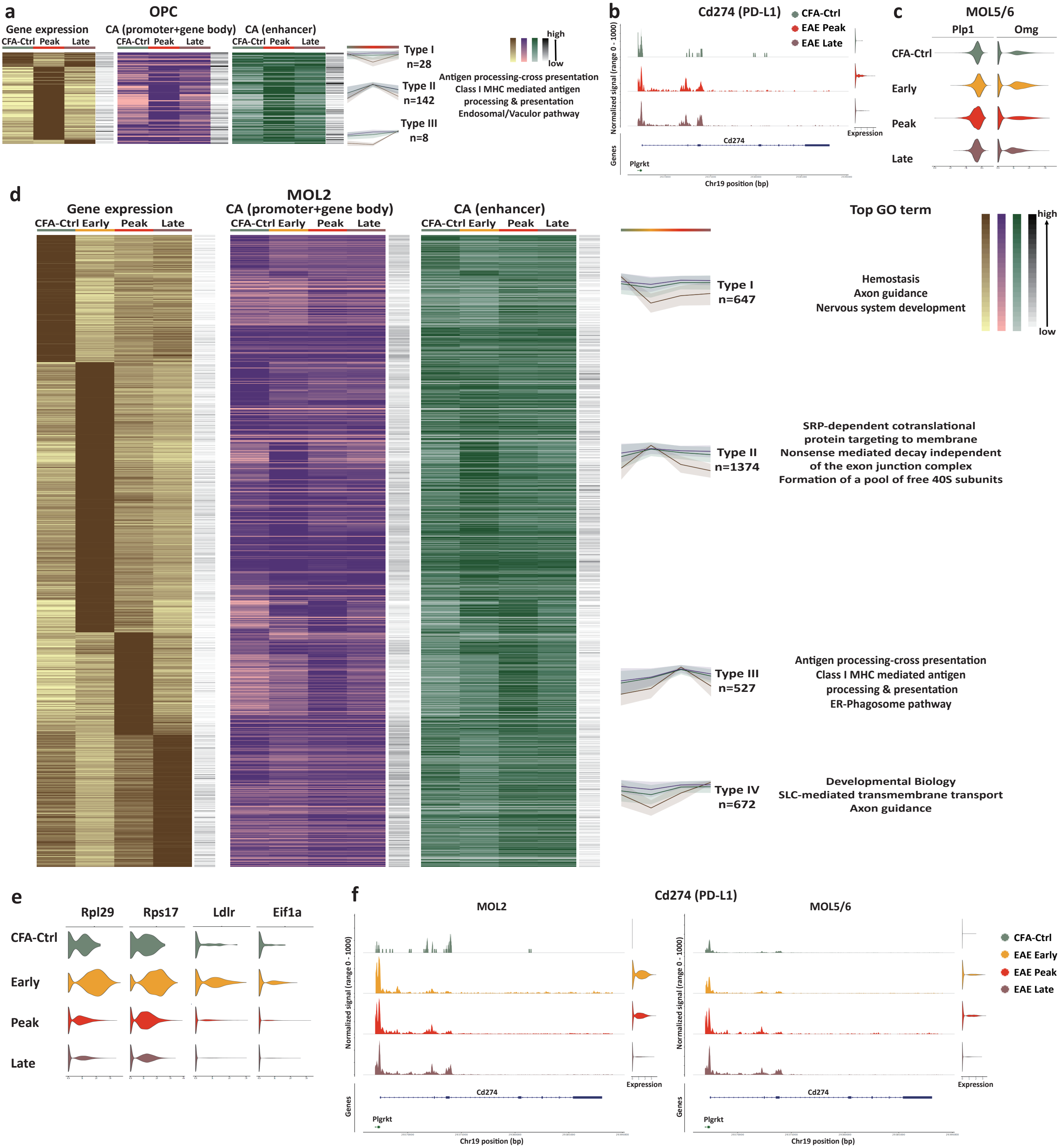


**Extended Data Fig. 5 Increased immune related transcription in OPCs and MOL2 at early and peak stages of EAE.**
a, Heatmaps of differentially expressed genes (in brown) between different time points in OPC, the chromatin accessibility at promoter and gene body (in purple) and at enhancer regions (in green) of the same gene (left), and the top GO terms of type II genes (middle). The black column on the side represents the gene raw counts. The line plots represent the mean of gene expression (line in brown), the chromatin accessibility at promoter and gene body (line in purple), and chromatin accessibility at enhancer regions (line in green) of genes in different groups (right).
b, Normalized tracks of chromatin accessibility (left) and expression (right) of Cd274 (PD-L1) in OPC.
c, Violin plots showing the expression of myelination related genes at different stages in MOL5/6.
d, Heatmaps of differentially expressed genes (in brown) between different time points in MOL5/6, chromatin accessibility at promoter and gene body (in purple) and at enhancer regions (in green) of the same gene (left), and the top GO terms of each group of genes (middle). The black column on the side represents the gene raw counts. The line plots represent the mean of gene expression (line in brown), the chromatin accessibility at promoter and gene body (line in purple), and chromatin accessibility at the enhancer regions (line in green) of genes in different groups (right).
e, Violin plot showing the expression of Rpl29, Rps17, Ldlr, and Eif1a in MOL2.
f, Normalized tracks of chromatin accessibility (left) and expression (right) of Cd274 (PD-L1) in MOL2 (left) and MOL5/6 (right).

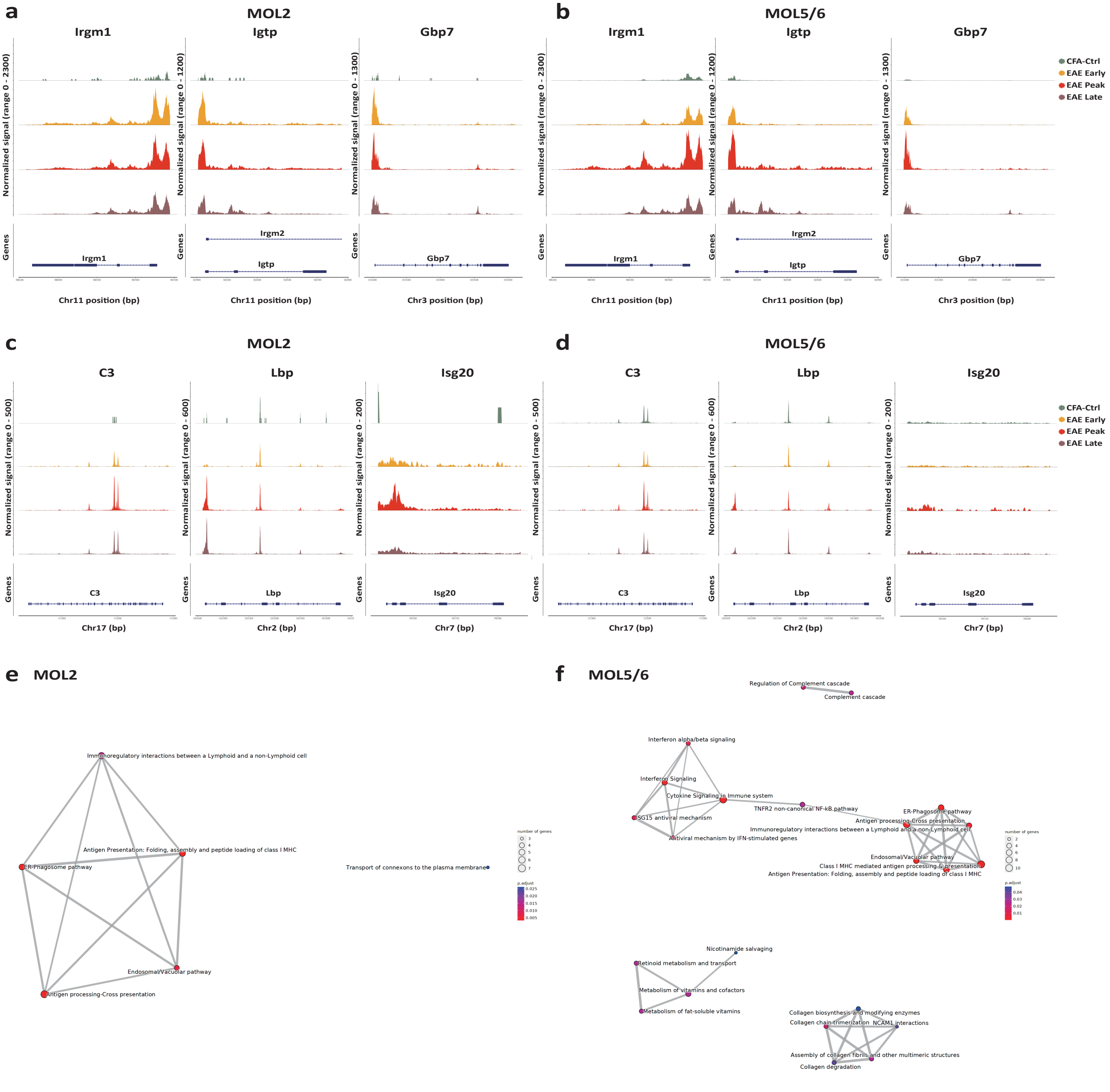


**Extended Data Fig. 6 Genes with significant changes in expression and chromatin accessibility among different disease stages were immune related.**
**a,b, Chromatin accessibility normalized tracks of representative immune related genes with significantly increase at peak stage in MOL2 (a) and MOL5/6 (b).**
**c,d, Normalized chromatin accessibility of C3, Lbp and Isg20 in MOL2 (c) and MOL5/6 (d) at each time point.**
**e,f, Network showing connections between enriched GO terms of genes with both differential expression and differential chromatin accessibility among different disease stages in MOL2 (e) and MOL5/6 (f).**

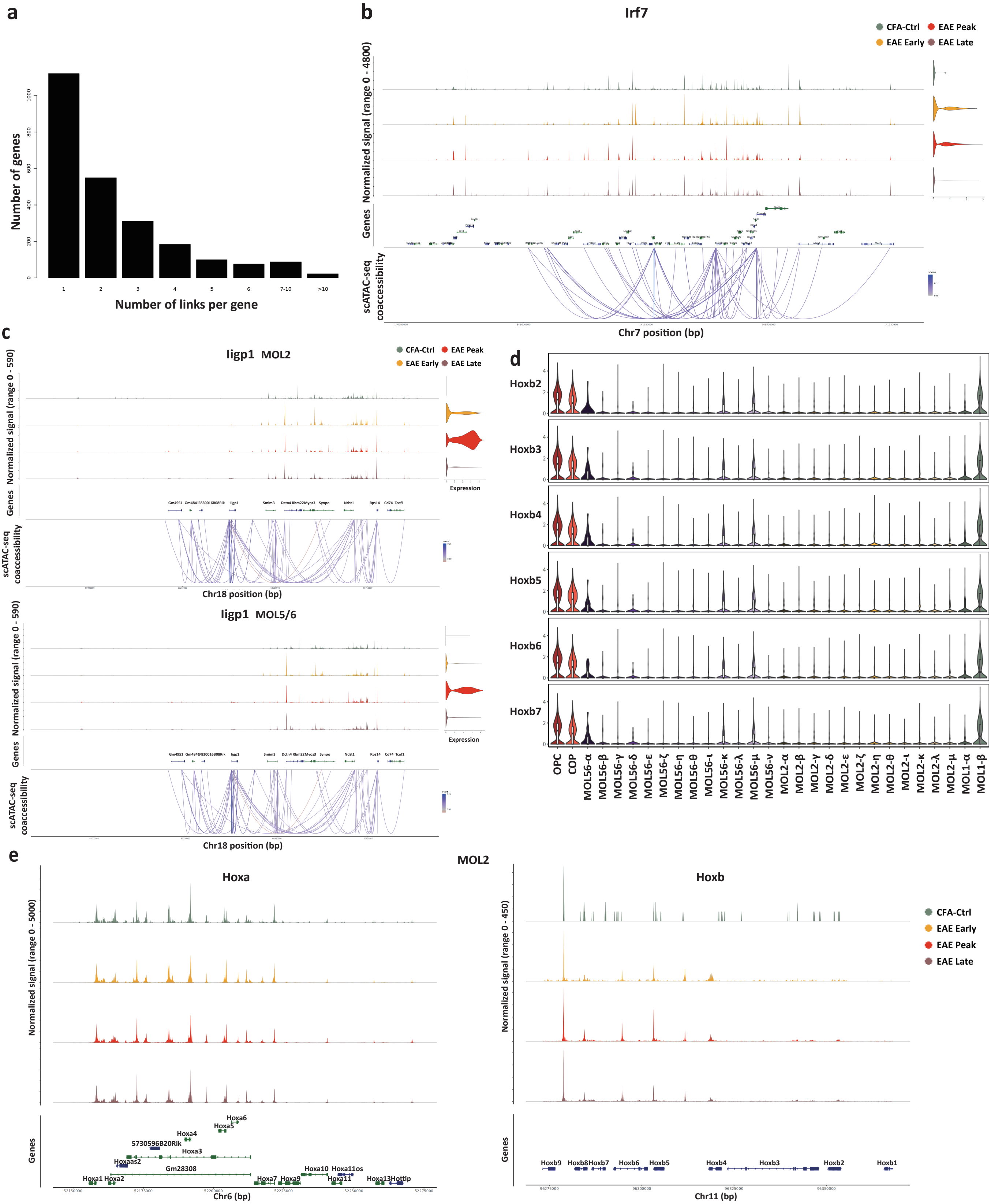


**Extended Data Fig. 7 Priming DORCs of immune related genes in MOL2 at early and peak stages and MOL5/6 at peak stage.**
**a**, The number of significant peak-gene associations for all genes.
**b**, Representative gene in Type II DORC in MOL2. The genomic track represents the accessibility of Irf7 at different time points, the links denote the significant correlation (pvalue < 0.05) between peaks and Irf7 (± 500 kb from TSSs). The violin plot shows the Irf7 expression in MOL2.
**c**, Representative gene in Type III DORC in MOL2 (left) and MOL5/6 (right). The genomic track represents the accessibility of Iigp1 at different time points, the links denote the significant correlation (pvalue < 0.05) between peaks and Iigp1 (± 500 kb from TSSs). The violin plot shows Iigp1 expression in MOL2 and MOL5/6.
**d**, DORCs score of Hoxb genes in OLG sub-cell types.
**e**, Normalized chromatin accessibility of Hoxa (left) and Hoxb (right) genes in MOL2.

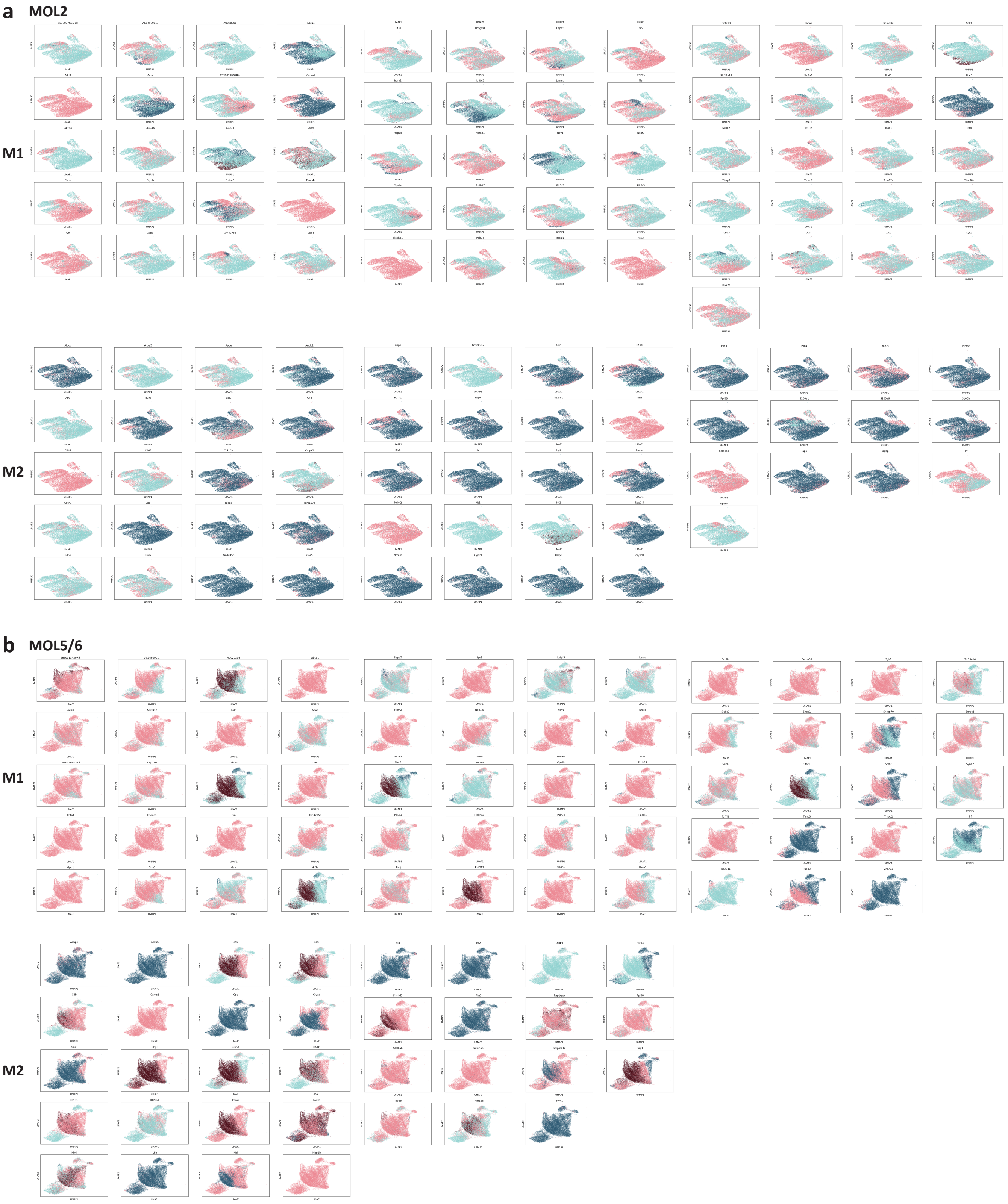


**Extended Data Fig. 8 MultiVelo analysis classified genes into two models.**
**a,b, UMAP of MOL2 (a) and MOL5/6 (b) colored by gene state assigned by MultiVelo, model 1 (M1, chromatin is closing before transcriptional repression) and model 2 (M2, chromatin is closing after transcriptional repression).**

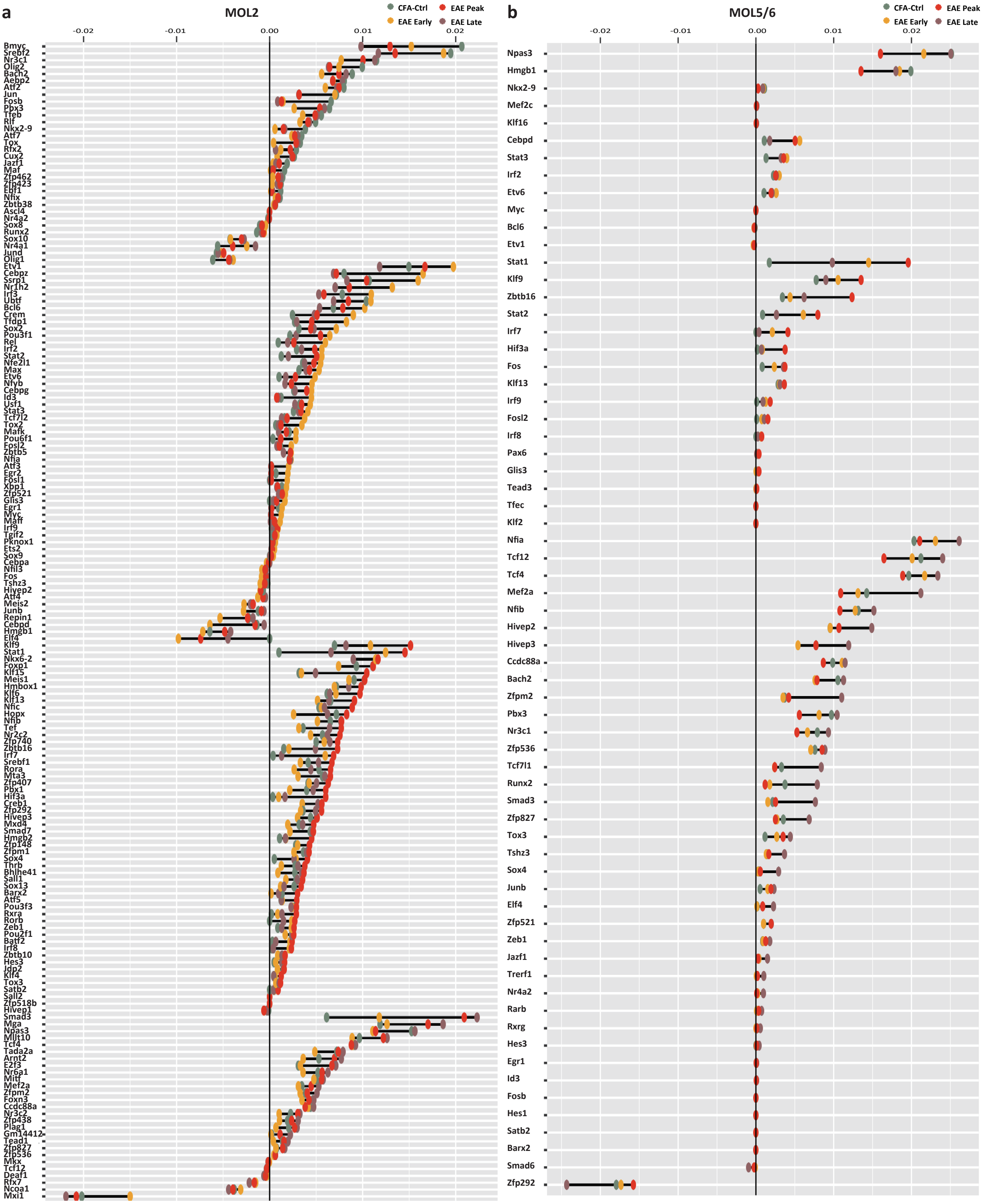


**Extended Data Fig. 9 Transcription factor activity**
a,b, Activity score of selected transcription factor at different stages in MOL2 (a) and MOL5/6 (b). x axis: transcription factor activity score, y axis: transcription factors grouped according to the time point at which transcription factor activity is highest.

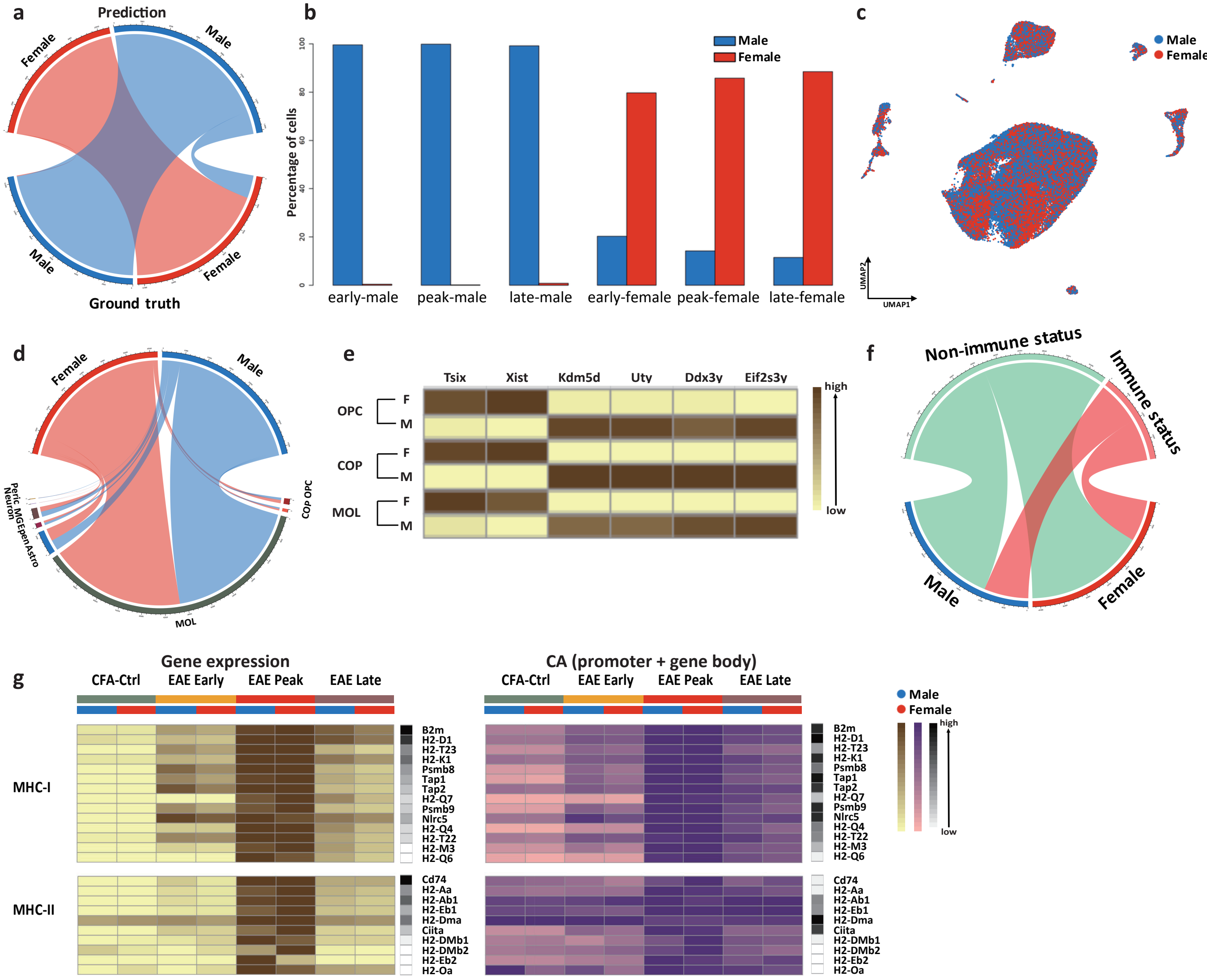


**Extended Data Fig. 10 No major differences were found in OLG between male and female regarding immune related genes.**
**a, Circos plot showing the prediction result (upper semicircle) and downsampled ground truth (bottom semicircle) of the sex.**
**b, Bar plot showing the percentage of cells predicted to be male (blue) or female (red) in the samples used to build the sex prediction model. These samples came from different stages of EAE and only contained cells from one gender.**
**c, Cell sex prediction on top of UMAP.**
**d, Circos plot showing the proportion of male and female cells in each cell type, downsampled by sex.**
**e, Heatmap showing the expression of differentially expressed genes between male and female in OPC, COP, MOL2, MOL5/6.**
**f, Number of cells with or without immune status for male and female (downsampled by sex).**
**g, Heatmaps of the expression (left) and chromatin accessibility (right) of MHC-I and -II genes in male and female at different stages in OLG. The black column on the side represents the gene raw counts.**

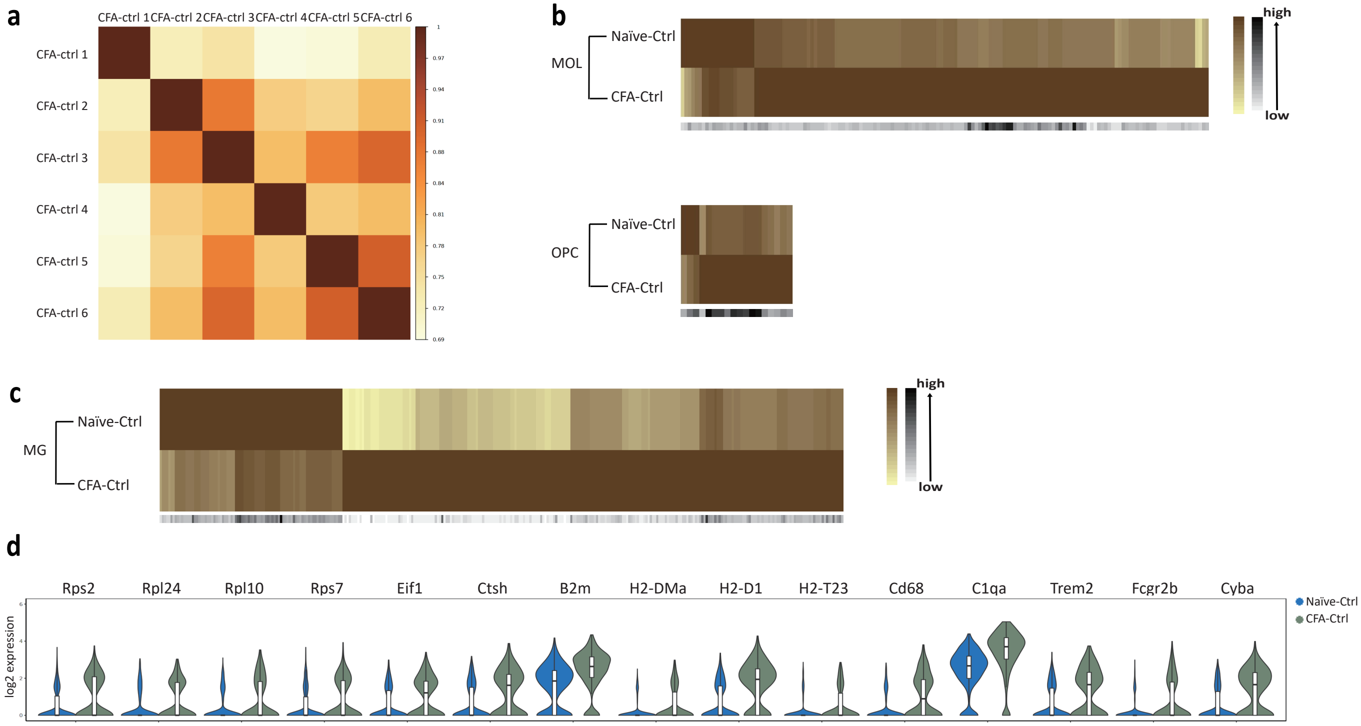


Extended Data Fig. 11 Immune genes were more enriched in MG from CFA-Ctrl than naïve-Ctrl.
a, Correlation matrix showing the Pearson correlation scores of MG between different CFA-Ctrl replicates.
b,c, Heatmaps showing normalized and scaled expression of differentially expressed genes between naïve-ctrl and CFA-Ctrl in OLG (b) and MG (c). The black raw represents the gene raw counts.
d, Violin plots showing some translation, MG activation, and immune system process genes which were significantly increased in CFA-Ctrl.
